# Semi-conservative transmission of DNA N^6^-adenine methylation in a unicellular eukaryote

**DOI:** 10.1101/2023.02.15.468708

**Authors:** Yalan Sheng, Yuanyuan Wang, Wentao Yang, Xue Qing Wang, Jiuwei Lu, Bo Pan, Bei Nan, Yongqiang Liu, Chun Li, Jikui Song, Yali Dou, Shan Gao, Yifan Liu

## Abstract

While DNA N^6^-adenine methylation (6mA) is best known in prokaryotes, its presence in eukaryotes has generated great interest recently. Biochemical and genetic evidence supports that AMT1, a MT-A70 family methyltransferase (MTase), is crucial for 6mA deposition in unicellular eukaryotes. Nonetheless, 6mA transmission mechanism remains to be elucidated. Taking advantage of Single Molecule Real-Time Circular Consensus Sequencing (SMRT CCS), here we provide definitive evidence for semi-conservative transmission of 6mA, showcased in the unicellular eukaryote *Tetrahymena thermophila*. In wildtype (WT) cells, 6mA occurs at the self-complementary ApT dinucleotide, mostly in full methylation (full-6mApT); hemi-methylation (hemi-6mApT) is transiently present on the parental strand of newly replicated DNA. In Δ*AMT1* cells, 6mA predominantly occurs as hemi-6mApT. Hemi-to-full conversion in WT cells is fast, robust, and likely processive, while *de novo* 6mA deposition in Δ*AMT1* cells is slow and sporadic. In *Tetrahymena*, regularly spaced 6mA clusters coincide with linker DNA of the canonical nucleosome arrays in the gene body. Importantly, *in vitro* methylation of human chromatin by reconstituted AMT1 complex recapitulates preferential targeting of hemi-6mApT sites in linker DNA, supporting AMT1’s intrinsic and autonomous role in maintenance methylation. We conclude that 6mA is transmitted by a semi-conservative mechanism: full-6mApT is split by DNA replication into hemi-6mApT, which is restored to full-6mApT by AMT1-dependent maintenance methylation. Our study dissects AMT1-dependent maintenance methylation and AMT1-independent *de novo* methylation, reveals a molecular pathway for 6mA transmission with striking similarity to 5-methyl cytosine (5mC) transmission at the CpG dinucleotide, and establishes 6mA as a *bona fide* eukaryotic epigenetic mark.

## Background

As a base modification, N^6^-adenine methylation can occur in both RNA (referred to as m6A) and DNA (6mA). m6A is ubiquitously present in rRNA, tRNA, and mRNA [1-4]. 6mA is extensively characterized in prokaryotes, involved in host genome defense, mismatch repair, and replication/transcription regulation [5]. 6mA in eukaryotes has also long been known, but its widespread presence is only lately realized [6-9]. 6mA studies in eukaryotes are complicated by varying abundance and divergent functions across species. In protists, green algae, and basal fungi, 6mA is abundant, enriched at the ApT dinucleotide, and associated with genes, all of which are consistent with its role as an epigenetic mark [7-12]. In animals, plants, and higher fungi, 6mA is scarce, promiscuous in its sequence context, and associated with silenced genomic regions [13-21]. In these organisms it remains controversial whether 6mA is an enzymatically deposited epigenetic mark, or merely a form of DNA damage, as the modified base, being a byproduct of RNA metabolism, is mis-incorporated into DNA [6, 22-25].

MT-A70 family of methyltransferases (MTases) are involved in N^6^-adenine methylation in eukaryotes [26, 27]. They are classified into several clades with distinct structures and functions [11, 26]. Two clades are represented by METTL3 and METTL14, forming a heterodimer for depositing m6A in mRNA [28]. As founding members of the family, METTL3 and METTL14 homologues are widely distributed and highly conserved in eukaryotes. Two additional clades are represented by AMT1 (also known as MTA1) and AMT6/7 (MTA9-B/MTA9), which are part of the eukaryotic 6mA MTase complex first identified in the protist *Tetrahymena thermophila* [10, 11]. METTL4/DAMT-1 are members of another clade [13], but they lack the DPPW motif critical for catalysis and their status as *bona fide* 6mA MTases is still not supported by biochemical evidence [10, 11, 26, 27]. Critically, AMT1 and AMT6/7 homologues are only found in protists, green algae, and basal fungi, while METTL4/DAMT-1 homologues are mostly found in animals, plants, and higher fungi [10, 11]. Phylogenetic distributions of these two deep branches of MT-A70 family members therefore closely match that of the two alternative modes of 6mA in eukaryotes [10, 11]. However, even in the best characterized *Tetrahymena* system, molecular mechanisms of 6mA transmission still need to be elucidated.

*Tetrahymena thermophila*, a ciliated protist, is the first eukaryote with 6mA identified in its nuclear DNA almost 50 years ago [29], and more recently, with AMT1, the eukaryotic 6mA-specific MTase, identified and characterized [10, 11]. *Tetrahymena* has been extensively studied as a model system for epigenetics and chromatin biology [30], and more specifically, for 6mA [7, 10-12, 29, 31, 32]. *Tetrahymena* contains within the same cytoplasmic compartment two types of nuclei, the somatic macronucleus (MAC) and the germline micronucleus (MIC) [33]. As the MAC is differentiated from the MIC, most transposable elements and repetitive sequences are first packaged into heterochromatin and subsequently removed in a RNAi and Polycomb-dependent pathway [34, 35]. While missing in the transcriptionally silent MIC, 6mA is abundantly present in the transcriptionally active MAC and associated with RNA polymerase II-transcribed genes, consistent with its role as a euchromatic mark [7, 29].

6mA is readily detected by Single Molecule Real-Time (SMRT) sequencing via its perturbation to DNA polymerase kinetics—specifically increase in the time between nucleotide incorporation, referred to as the inter-pulse duration (IPD) (Figure 1A) [36, 37]. Genome-wide mapping of 6mA in eukaryotes has previously been achieved only at the ensemble level, by combining different DNA molecules covering the same genomic position to overcome random fluctuations in IPD—an approach referred to as Continuous Long Reads (CLR) [7, 10, 11]. Effective implementation of Circular Consensus Sequencing (CCS; also known as PacBio HiFi Sequencing), by combining reads from multiple passes of the same DNA template (Figure 1B) [22, 36-38], allows us to accurately map 6mA distribution in the *Tetrahymena* MAC genome at the single molecule level and rigorously establish AMT1-dependent semi-conservative transmission of 6mA.

**Figure 1.**
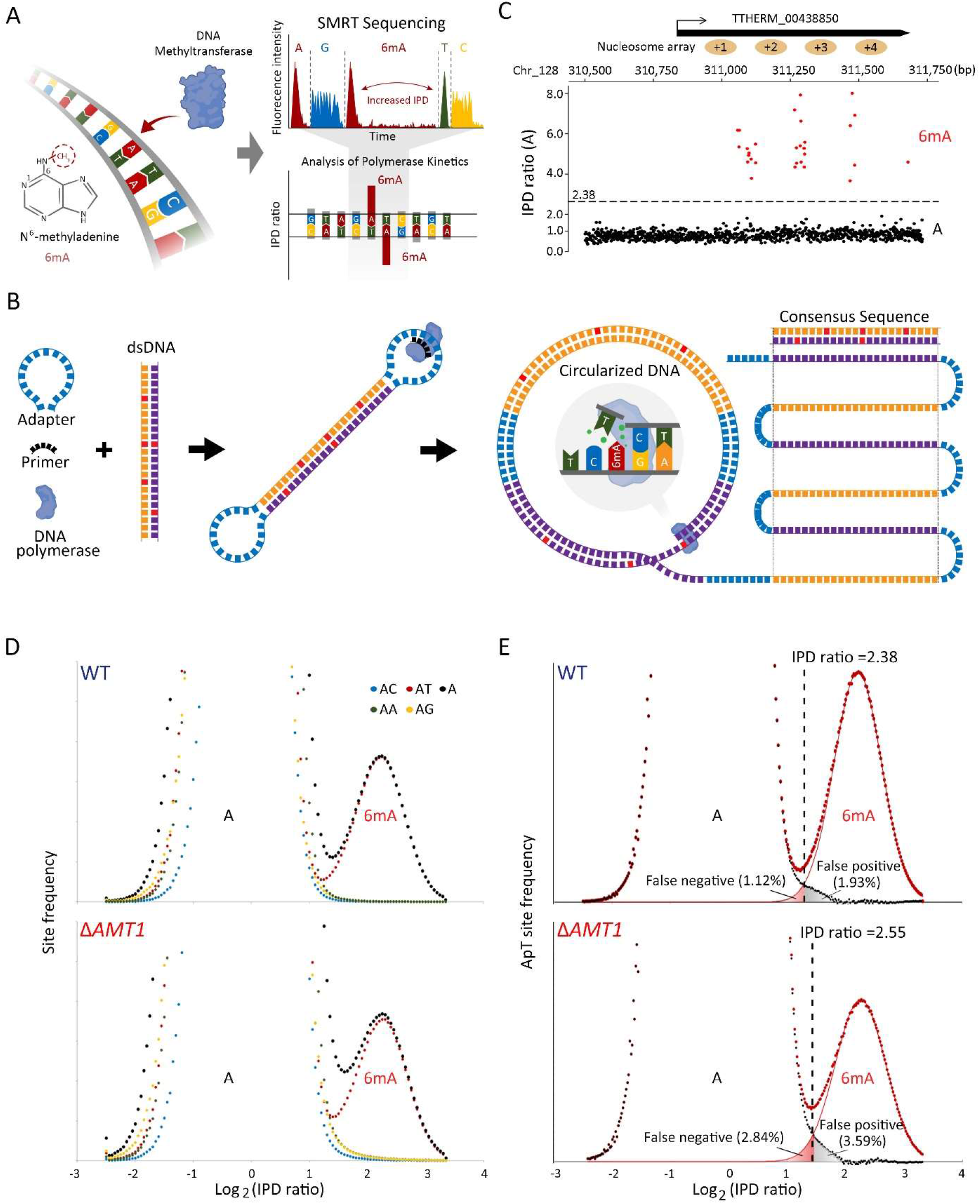
6mA detection by SMRT CCS. A. Schematic for 6mA detection by SMRT sequencing. B. Schematic for SMRT CCS. C. IPD ratios (IPDr) for all A sites in a typical SMRT CCS read mapped to the *Tetrahymena* MAC reference genome. The IPDr threshold was set at 2.38, separating 6mA from unmodified A. Note the localization of 6mA clusters in linker DNA between the canonical nucleosome array within the gene body. D. IPDr distribution (log2) of all A sites in WT (top) and Δ*AMT1* cells (bottom). Also plotted were distributions for A sites at the ApA, ApC, ApG, and ApT dinucleotide, respectively. E. Deconvolution of the 6mA peak and the unmodified A peak for IPDr distributions (log2) at the ApT dinucleotide. Note the low false positive and false negative rates of 6mA calling in WT (top) and Δ*AMT1* cells (bottom).

## Results

### 6mA detection at the single molecule level

We developed a SMRT CCS-based pipeline to map 6mA on individual DNA molecules from *Tetrahymena* (Figure S1A). As 6mA calling accuracy scaled with the number of passes for CCS, we set a stringent threshold (≥30×) for high-quality reads (Figure S1B). We used the CCS read of a DNA molecule as its own reference sequence in IPD analysis, yielding averaged and standardized IPD ratios (IPDr) for each site, relative to the *in silico* reference for its unmodified counterpart (Figure S1A) [22]. A typical DNA molecule from wildtype (WT) *Tetrahymena* cells showed low IPDr for most adenine (A) sites and a few clusters with high IPDr (Figure 1C). As most A sites are presumably unmodified, they formed a baseline of IPDr around 1, with low dispersion across the read length (Figure 1C). As exceptions, we found DNA molecules with global anomalies in IPDr, whose baseline dispersed or deviated from 1 (Figure S1C, D), possibly due to a compromised DNA polymerase. We also found DNA molecules with local anomalies in IPDr, which contained one or more clusters of high IPDr G/C/T as well as A sites (Figure S1E), attributable to DNA damage (*38*). Both exceptions were removed from further analysis.

We next mapped CCS reads back to the *Tetrahymena* MAC, MIC, and mitochondrion reference genomes (Figure S2A-D). Most were aligned across the entire read to a single genomic locus (Figure S2B). There were some chimeric reads with different parts aligned to separate genomic loci (Figure S2B), attributable to concatenation during sequencing library preparation. Their constituent DNA molecules were resolved before further analysis (Figure S2C, D). For DNA molecules fully mapped to the MAC genome, their IPDr for A sites exhibited a bimodal distribution: a large peak with low IPDr corresponding to unmodified A and a small peak with high IPDr corresponding to 6mA (Figure 1D: top). A similar bimodal distribution was observed when we focused on A sites within the ApT dinucleotide (Figure 1D: top). The 6mA peaks of these two distributions were almost superimposable (Figure 1D: top, Figure S2E: left). In contrast, IPDr distributions of A sites within the ApA/ApC/ApG dinucleotides all exhibited a single peak with low IPDr (Figure 1D: top). These analyses indicate that 6mA is exclusively associated with the ApT dinucleotide 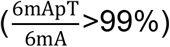. Our conclusion disagrees with previous estimate of substantial 6mA in non-ApT dinucleotides (12%) based on SMRT CLR [10, 11].

We deconvoluted the 6mA peak and unmodified A peak in the IPDr distribution for the ApT dinucleotide: the 6mA peak was closely fitted by a Gaussian distribution curve, while the unmodified A peak was deduced as the differential between the original data and the Gaussian fit (Figure 1E: top). We set the threshold for 6mA calling at the intersection of the two peaks (IPDr=2.38) and estimated that the false positive and false negative rates of 6mApT calling were 1.93% and 1.12%, respectively (Figure 1E: top). Note that CCS and CLR results converge at genomic positions of high 6mA coverage (where CLR can also make high confidence calls) but diverge substantially at low 6mA coverage (Figure S3). We calculated that 6mApT represented 1.86% of all ApT sites in DNA molecules fully mapped to the MAC genome. Using the same threshold, 6mApT was called only at low levels for DNA molecules specifically mapped to the MIC (0.017%) or mitochondrion (0.014%) (Figure S2D, Table S1). 6mA at the ApC/ApG/ApA dinucleotides was called at low levels regardless of their mapping (Table S1). These low level 6mA calls probably represent the background noise. We conclude that 6mA occurs exclusively at the ApT dinucleotide in the MAC.

### Distinguishing four methylation states of ApT duplexes

SMRT CCS makes strand-specific 6mA calls, as the DNA polymerase alternately passes through the Watson strand (W, defined as the forward strand in the reference genome) and the Crick strand (C, reverse) of a DNA template (Figure 1B) [36]. For self-complementary ApT duplexes, we plotted their distribution according to IPDr values of A sites on W and C, respectively, and found four groups with diagonal symmetry, corresponding to four methylation states: full methylation, methylation only on W (hemi-W), methylation only on C (hemi-C), and no methylation (Figure 2A-C). We set two thresholds to demarcate these four groups. For bulk ApT sites, the high IPDr threshold set for 6mA calling (IPDr=2.38) was in large part to compensate for the predominance of the unmodified A peak over the 6mA peak (98% vs. 2%). However, for an ApT site reverse complementary to a 6mApT site in ApT duplexes, the unmodified A peak was instead dominated by the 6mA peak; this caused the 6mA calling threshold to shift substantially to the left (IPDr=1.57) (Figure 2C: left). Based on this demarcation, we estimated that 89% methylated ApT duplexes were full methylation (full-6mApT), while 11% were hemi-methylation (hemi-6mApT) (Table 1, Figure 2C: left, Figure 2D: top). Importantly, consistent evaluation of the full- and hemi-6mApT percentages was obtained in duplicate experiments (Figure S4). Our results establish the predominance of full-6mApT over hemi-6mApT in WT *Tetrahymena* cells, in contrast to the near parity assessment (54% and 46%, respectively) based on CLR [11]. Note that only SMRT CCS can distinguish between hemi- and full-6mApT at the single molecule level, while CLR must extrapolate from the ensemble level.

**Table 1.**
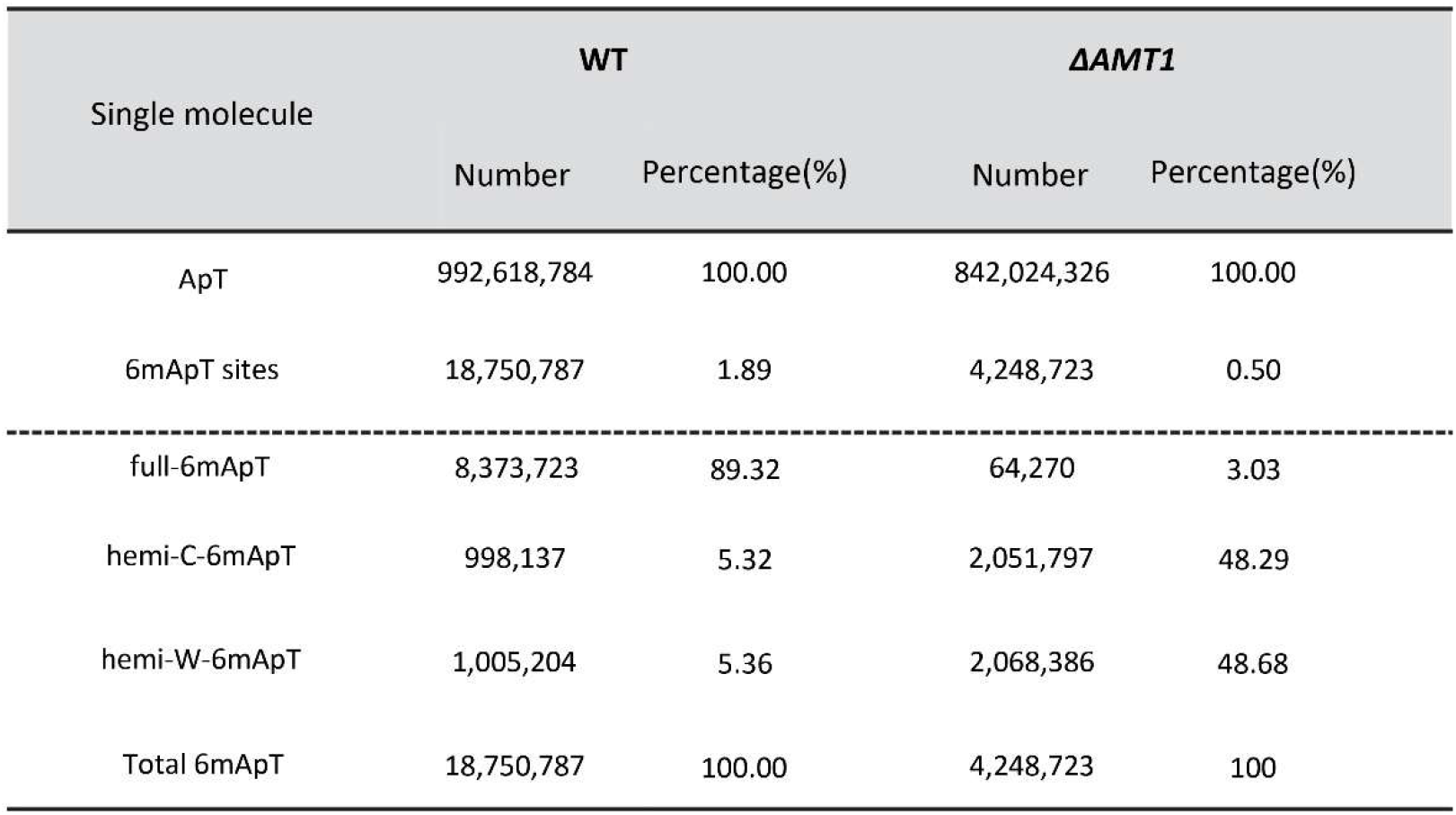
6mA statistics in WT and Δ*AMT1* cells. Top: the number of total ApT (with or without modification) and 6mApT sites in DNA molecules fully mapped to the MAC. Both W and C are counted. Percentage of DNA methylation is also calculated 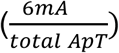. Bottom: the number of full-6mApT and hemi-6mApT duplexes. Note that each full-6mApT duplex contains two 6mA sites, while each hemi-6mApT duplex only contains one site. Percentages of full-6mApT and hemi-6mApT duplexes are also calculated.

| Single molecule | WT |  | *ΔAMT1* |  |
| --- | --- | --- | --- | --- |
|  | Number | Percentage(%) | Number | Percentage(%) |
| ApT | 992,618,784 | 100.00 | 842,024,326 | 100.00 |
| 6mApT sites | 18,750,787 | 1.89 | 4,248,723 | 0.50 |
| full-6mApT | 8,373,723 | 89.32 | 64,270 | 3.03 |
| hemi-C-6mApT | 998,137 | 5.32 | 2,051,797 | 48.29 |
| hemi-W-6mApT | 1,005,204 | 5.36 | 2,068,386 | 48.68 |
| Total 6mApT | 18,750,787 | 100.00 | 4,248,723 | 100 |

**Figure 2.**
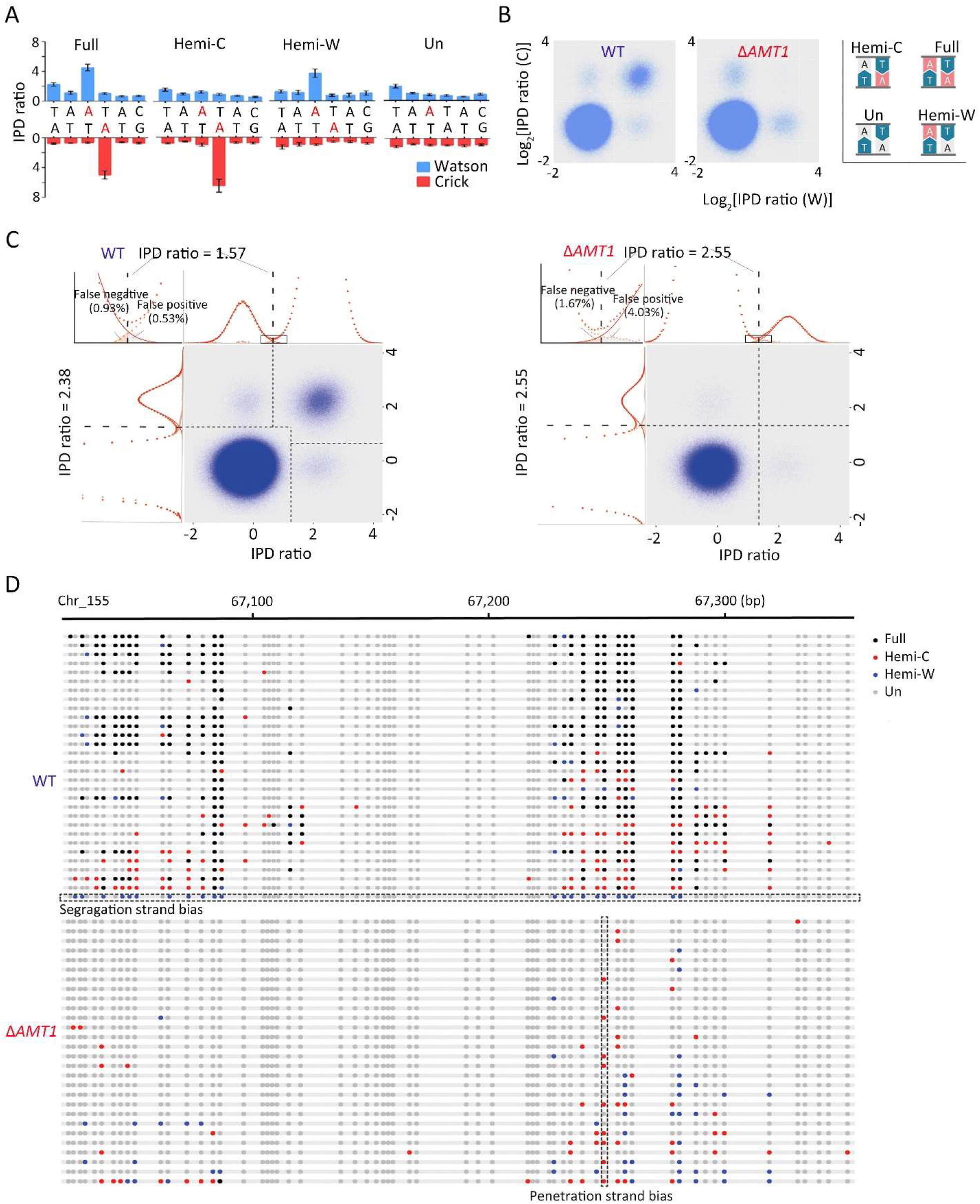
Distinguishing four methylation states of ApT duplexes. A. Four states of ApT duplexes: full methylation, hemi-W, hemi-C, and unmethylated, distinguished by IPDr of adenine sites on W and C, respectively. B. Distribution of ApT duplexes according to IPDr of adenine sites on W and C, respectively. Note the abundance of the full methylation state in WT and its absence in Δ*AMT1* cells. C. Demarcation of the four methylation states of ApT duplexes in WT (left) and Δ*AMT1* cells (right) by their IPDr on W and C, respectively. Left: For bulk ApT duplexes, the IPDr threshold for 6mA calling was set at 2.38, according to deconvolution based on Gaussian fitting of the small 6mA peak. For ApT duplexes with one 6mA as defined above, the IPDr threshold for calling 6mA on the opposite strand was set at 1.57, according to deconvolution based on Gaussian fitting of the small unmodified A peak. Right: For bulk ApT duplexes, the IPDr threshold for 6mA calling was set at 2.55, according to deconvolution based on Gaussian fitting of the small 6mA peak. For ApT duplexes with one 6mA as defined above, the IPDr threshold for calling 6mA on the opposite strand was also set at 2.55, according to deconvolution based on Gaussian fitting of the small 6mA peak. D. Typical DNA molecules from *Tetrahymena* WT (top) and Δ*AMT1* cells (bottom). Note ApT duplexes with distinct methylation states (colored dot) distributed along individual DNA molecules (gray line). A DNA molecule with strong segregation strand bias in WT cells and a genomic position with strong penetration strand bias in Δ*AMT1* cells were marked.

We define 6mA penetration for each genomic position as the ratio between the number of 6mA sites and all adenine sites (with or without modification) in all SMRT CCS reads. For WT cells, 6mA penetration for most ApT positions showed no significant bias for either W or C (Figure 6D: top); with increasing sequencing coverage, 6mA penetration from both strands tended to converge (Figure 6D: middle). In other words, at the ensemble level, most ApT positions in the genome were methylated at similar levels on W or C. We did not observe biased 6mA penetration even in asymmetrically methylated ApT positions reported previously [11], and attributed them as a CLR artefact. Our result is consistent with DNA replication splitting a full-6mApT into a hemi-W and a hemi-C.

### Segregation of hemi-6mApT to the old strand after DNA replication

We next investigated segregation of hemi-W and hemi-C at the single molecule level. We focused on DNA molecules with multiple hemi-6mApT, henceforth referred to as hemi^+^ molecules (Figure 2D: top, Figure S5). Their levels oscillated with cell cycle progression, starting low for cells synchronized at G1 phase, climbing to the peak for cells in S phase, and declining for post-replicative and dividing cells (Figure 3A). In the vast majority of hemi^+^ molecules, hemi-6mApT were not randomly distributed across both strands; instead, their constituent 6mA sites were segregated with a strong bias for one strand (Figure 3B, C and Figure S5B). These results all support hemi^+^ molecules as the product of DNA replication. We also noted that this segregation was not always absolute: a minority of hemi-6mApT were occasionally detected on the opposite strand (Figure S5B). This is most likely due to *de novo* methylation, either AMT1-dependent (see **AMT1-dependent maintenance methylation**) or AMT1-independent (see **AMT1-independent *de novo* methylation**).

**Figure 3.**
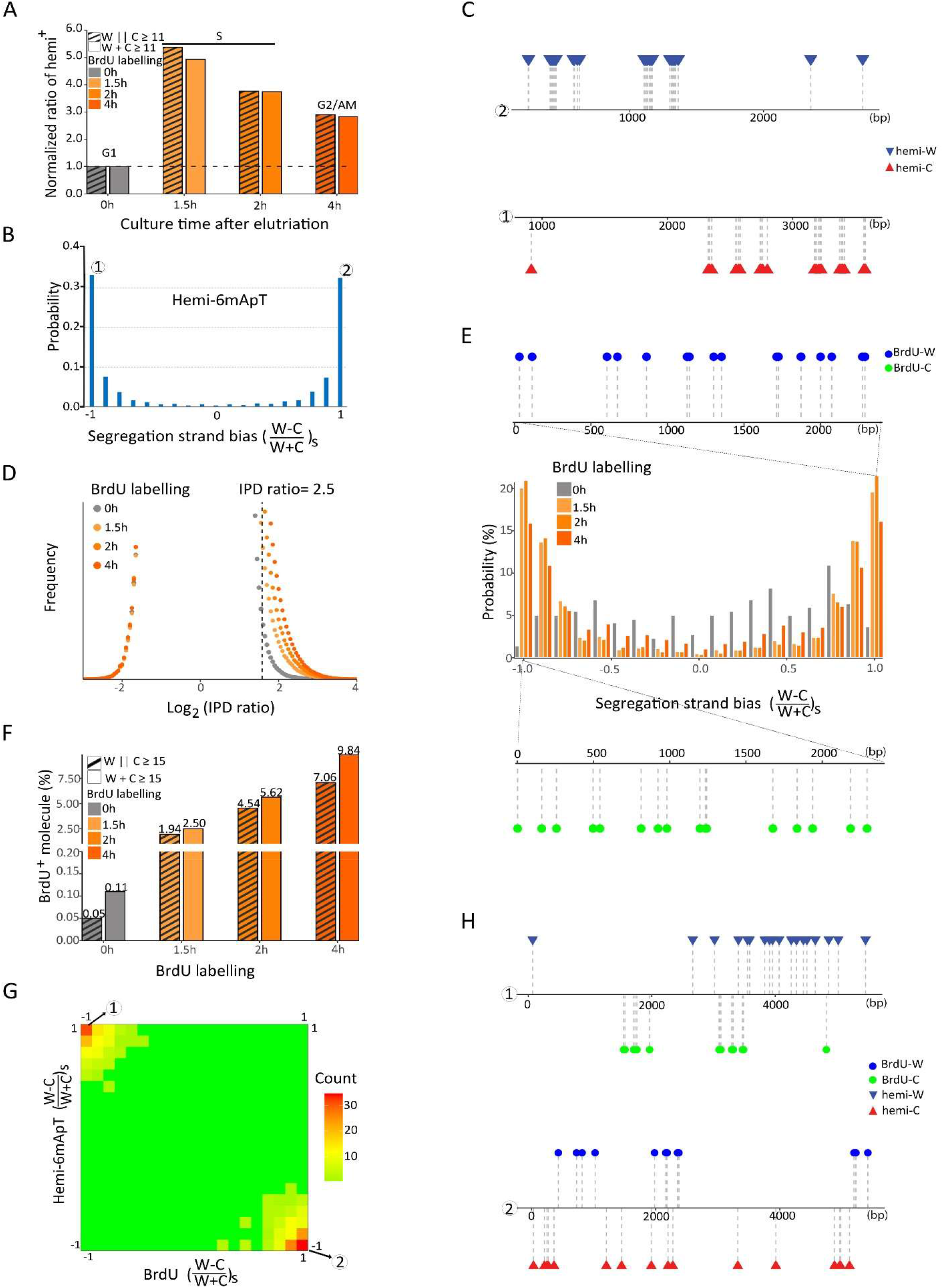
Segregation of hemi-6mApT to the old strand after DNA replication. A. Hemi^+^ molecules are enriched in S phase. *Tetrahymena* cells were synchronized at G1 phase by centrifugal elutriation and released for growth in the fresh medium [55]. Four time points were taken (0, 1.5, 2, and 4h after release) for SMRT CCS. Hemi^+^ molecules were defined as DNA molecules with a total count of more than 11 hemi sites (W+C≥11) or with more than 11 hemi sites on one strand (W||C≥11). The count of hemi^+^ molecules was normalized first against the counts of total DNA molecules and the 0h (G1) value. B. Hemi-6mApT sites in hemi^+^ molecules exhibit strong segregation strand bias. Segregation strand bias for hemi-6mApT is defined as the difference-sum ratio between hemi-W and hemi-C: 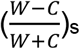. C. Typical DNA molecules with hemi-6mApT fully segregated to W or C, corresponding to segregation strand bias of +1 and -1, as marked in Figure 3B. D. IPDr distribution of T sites in genomic DNA samples of synchronized *Tetrahymena* cells with BrdU-labeling (1.5h, 2h, and 4h) or without (0h). The IPDr threshold for calling BrdU was set at 2.5. E. BrdU sites in BrdU^+^ molecules exhibit strong segregation strand biases. BrdU^+^ molecules were defined as DNA molecules with a total count of more than 15 BrdU sites (W+C≥15). Segregation strand bias for BrdU was defined as the difference-sum ratio between BrdU sites on W and C: 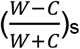. Also shown are typical BrdU^+^ molecules with BrdU fully segregated to W or C, corresponding to segregation strand bias of +1 and -1, respectively. F. Correlation between BrdU-labeling and BrdU^+^ molecules. BrdU^+^ molecules were alternatively defined as DNA molecules with a total count of more than 15 BrdU sites (W+C≥15), or with more than 15 BrdU sites on one strand (W||C≥15). The latter is more stringent than the former, as it is more selective for DNA molecules with strong strand segregation bias. G. Hemi-6mApT and BrdU are segregated to opposite strands of the DNA duplex. Distribution of hemi^+^/BrdU^+^ molecules (hemi-6mApT: W||C≥11; BrdU: W||C≥15) according to their segregation strand bias for hemi-6mApT and BrdU, respectively. H. Typical hemi^+^/BrdU^+^ molecules with hemi-6mApT and BrdU fully segregated to opposite strands, corresponding to segregation strand bias of (−1, +1) and (+1, - 1), as marked in Figure 3G.

To determine whether hemi-6mApT were segregated to the old strand or the newly synthesized strand after DNA replication, we labeled *Tetrahymena* cells with 5-bromo-2’-deoxyuridine (BrdU) (Figure S6A). BrdU substitution of thymidine resulted in IPDr increase, readily detected by SMRT CCS (Figure 3D, Figure S6B). To eliminate interference from 6mA, we masked regions adjacent to 6mApT sites (both strands: -10 to +10bp) from further analysis (Figure S6B). To increase the chance to correctly identify BrdU-labeled DNA molecules, we focused on those with multiple BrdU calls, henceforth referred to as BrdU^+^ molecules (Figure 3E, F). In BrdU-labeled samples, BrdU sites in BrdU^+^ molecules were mostly segregated to one strand (Figure 3E). In the unlabeled sample, “BrdU” sites were more evenly distributed across both strands, consistent with miscalls due to random fluctuations in IPDr (Figure 3E). Our approach was further validated by strong correlations between BrdU labeling and BrdU^+^ molecules: ***1)*** there were many BrdU^+^ molecules in BrdU-labeled samples, but few in the unlabeled sample (the percentage was further reduced when focusing on BrdU^+^ molecules with strong biases in strand segregation); and ***2)*** the percentage of BrdU^+^ molecules increased progressively with longer labeling time (Figure 3F). BrdU segregation was often not absolute (Figure 3E), attributable to high false positive rates of BrdU calling (Figure 3D). Nonetheless, the large number of BrdU sites in BrdU^+^ molecules allow us to identify the newly synthesized DNA strand with high confidence.

There were significant overlaps between BrdU^+^ and hemi^+^ molecules in BrdU-labeled samples (Figure S6C). We focused on BrdU^+^/hemi^+^ double-positive molecules representing post-replicative DNA (Figure 3G, H). Critically, BrdU and hemi-6mApT always exhibited the opposite biases for strand segregation in BrdU^+^/hemi^+^ molecules (Figure 3G, H). This result indicates that after DNA replication, hemi-6mApT is essentially excluded from the newly synthesized strand and only associated with the old strand.

### AMT1-dependent maintenance methylation

To complete semi-conservative transmission of 6mA, hemi-6mApT needs to be restored to full-6mApT by maintenance methylation before next round of DNA replication. We investigated whether maintenance methylation was dependent on AMT1. In Δ*AMT1* cells, SMRT CCS showed that 6mA was still predominantly associated with the ApT dinucleotide 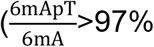; Figure 1D: bottom, Figure S2E: right), in contrast to our previous estimation of a majority of 6mA in non-ApT dinucleotides (53%) based on SMRT CLR [11]. While WT cells contained mostly full-6mApT (89%), there were few in Δ*AMT1* cells (3%) (Figure 1E: bottom, Figure 2B, Figure 2C: right, Figure 2D: bottom, Table 1). The predominant hemi-6mApT in Δ*AMT1* cells is presumably the product of a dedicated *de novo* MTase. We conclude that AMT1 is required for hemi-to-full conversion, *i*.*e*., maintenance methylation.

AMT1 is part of a multi-subunit MTase complex [10, 39]. We next reconstituted AMT1 complex comprising bacterially expressed AMT1, AMT7, AMTP1, and AMTP2 (also known as MTA1, MTA9, p1, and p2 [10]) (Figure 4A). Using a 12-bp DNA substrate with a single centrally located hemi-6mApT, we tested the reconstituted complex for *in vitro* methylation and evaluated its steady-state kinetics (Km=0.55μM, k_cat_=0.84min^-1^; Figure 4B). We also compared two 27-bp DNA substrates with the same primary sequence: the hemi-methylated substrate contained two 6mApT sites segregated to one strand, while its counterpart was completely unmodified (Figure 4C). The hemi-methylated substrate recorded 11.5× higher activity than the unmodified substrate (Figure 4C), a much more dramatic advantage than previously reported [10]. AMT1 complex therefore strongly prefers maintenance methylation to *de novo* methylation.

**Figure 4.**
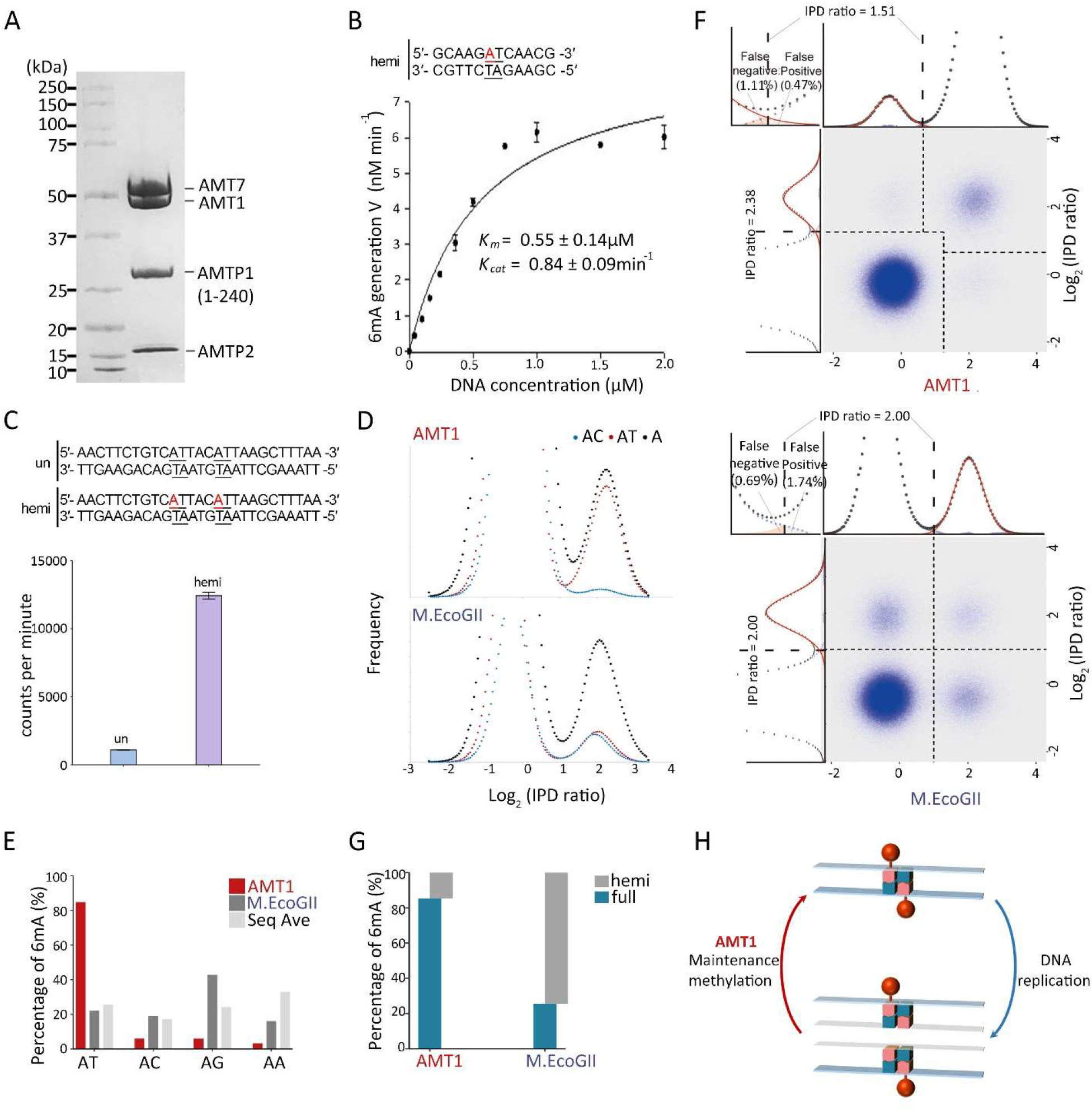
In vitro methyltransferase activity of AMT1 complex. A. SDS-PAGE of *in vitro* reconstituted AMT1 complex comprising AMT1, AMT7, AMTP1 (1-240 aa), and AMTP2. B. Steady-state kinetics of AMT1 complex on a hemi-methylated substrate (hemi). It contains a single ApT duplex (underlined), which is hemi-methylated (red). C. Methylation of the unmodified (un) and hemi-methylated (hemi) substrates. Both contain two ApT duplexes (underlined), which are either unmodified or hemi-methylated (red). D. IPDr distributions for total adenine, adenine in the ApT dinucleotide, and adenine in ApC dinucleotide, after *in vitro* methylation of human chromatin by either AMT1 complex (top) or M.EcoGII (bottom). E. 6mA distribution in all four ApN dinucleotides, after *in vitro* methylation of human chromatin by either AMT1 complex or M.EcoGII. ApN frequencies in SMRT CCS reads are also plotted for comparison. F. Demarcation of the four methylation states of ApT duplexes by their IPDr on W and C, in human chromatin methylated by AMT1 complex (top) or M.EcoGII (bottom). AMT1 complex methylation pattern is reminiscent of that in WT *Tetrahymena* cells, with strong preference for full-6mApT, as indicated by shift in the IPDr threshold for calling bulk 6mA or full-6mApT. M.EcoGII methylation pattern is reminiscent of that in Δ*AMT1* cells, with no preference for full-6mApT, as indicated by the same IPDr threshold for calling bulk 6mA or full-6mApT. G. Relative abundance of hemi-6mApT and full-6mApT in human chromatin methylated by either AMT1 complex or M.EcoGII. H. Model: semi-conservative transmission of 6mA in WT *Tetrahymena* cells.

We also performed *in vitro* methylation of human chromatin using the reconstituted AMT1 complex (Figure 4D-G, Figure S7); as a control, we used M.EcoGII, a prokaryotic MTase targeting adenine sites in any sequence context [40]. Due to scarcity of endogenous 6mA in human genomic DNA [6, 22, 41, 42], all 6mA sites revealed by SMRT CCS were essentially attributable to the added MTases. We found that 85% of 6mA sites were at the ApT dinucleotide after AMT1 complex treatment; only 22% of 6mA sites were so after M.EcoGII treatment, close to the ApT frequency in sequenced DNA molecules (Figure 4D, E, Figure S7C). The substantial 6mA in non-ApT dinucleotides (15%) after AMT1 complex treatment is consistent with previous characterization of 6mA-MTase activity partially purified from *Tetrahymena* [32]. Therefore, *in vitro* methylation catalyzed by AMT1 complex occurs preferentially at the ApT dinucleotide, but not exclusively, as *in vivo*.

Methylation of ApT sites was at similar levels and far from saturation in both AMT1 complex and M.EcoGII-treated samples 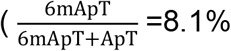 and 9.5%, respectively). However, 85% methylated ApT duplexes were full-6mApT after AMT1 treatment, while only 26% were so after M.EcoGII treatment (Figure 4F, G). In the case of AMT1 complex, we found that the IPDr threshold for calling 6mA in ApT duplexes with 6mA on the opposite strand was substantially shifted to the left, when compared with the IPDr threshold for calling 6mA in bulk ApT duplexes (conditional probability**≠**unconditional probability) (Figure 4F: top). Importantly, a very similar shift in the IPDr threshold for calling 6mA in ApT duplexes was observed in WT *Tetrahymena* cells (Figure 2C: left). In the case of M.EcoGII, the IPDr threshold for calling 6mA in ApT duplexes stayed the same, regardless of the methylation state of the opposite strand (conditional probability**=**unconditional probability) (Figure 4F: bottom). Therefore, M.EcoGII does not prefer maintenance methylation (hemi-to-full conversion) over *de novo* methylation (un-to-hemi conversion). As a corollary, full-6mApT is generated by random combination of two independent methylation events. In contrast, AMT1-dependent maintenance methylation is much faster than *de novo* methylation, leading to accumulation of full-6mApT and depletion of hemi-6mApT under *in vitro* as well as *in vivo* conditions. Therefore, preferential targeting of ApT, especially hemi-6mApT, is an intrinsic and autonomous property of AMT1 complex. We conclude that 6mA is transmitted by a semi-conservative mechanism in *Tetrahymena*: full-6mApT is split by DNA replication into hemi-6mApT, which is restored to full-6mApT by AMT1-dependent maintenance methylation (Figure 4H).

### Preferential methylation of linker DNA by AMT1 complex

Previous studies have shown that in unicellular eukaryotes, 6mA distribution is connected to nucleosome distribution, suggesting that 6mA transmission relies on the chromatin environment as well as the sequence context [7, 8, 10]. SMRT CCS revealed that on individual DNA molecules from WT *Tetrahymena* cells, 6mA sites generally distributed in clusters separated by regular intervals (Figure 5A). Autocorrelation analysis confirmed that 6mA sites were strongly phased at the single molecule level, oscillating with cycles of ∼200bp (Figure 5B). Furthermore, 6mA clusters from different DNA molecules were often coarsely aligned to the same genomic region (Figure 5A). Indeed, 6mA distribution was also phased at the ensemble level, just like nucleosome distribution in *Tetrahymena* (Figure 5C: top). Autocorrelation analysis showed that 6mA and nucleosome distributions in *Tetrahymena* shared the same cycle of ∼200bp (Figure 5C: top); cross-correlation analysis showed that 6mA and nucleosome distributions were offset by ∼100bp and in opposite phases (Figure 5C: bottom). We also found that 6mA peaks coincided with nucleosome troughs downstream of transcription start sites (Figure 5A, Figure S8A). Therefore, 6mA is preferentially associated with linker DNA in *Tetrahymena*.

**Figure 5.**
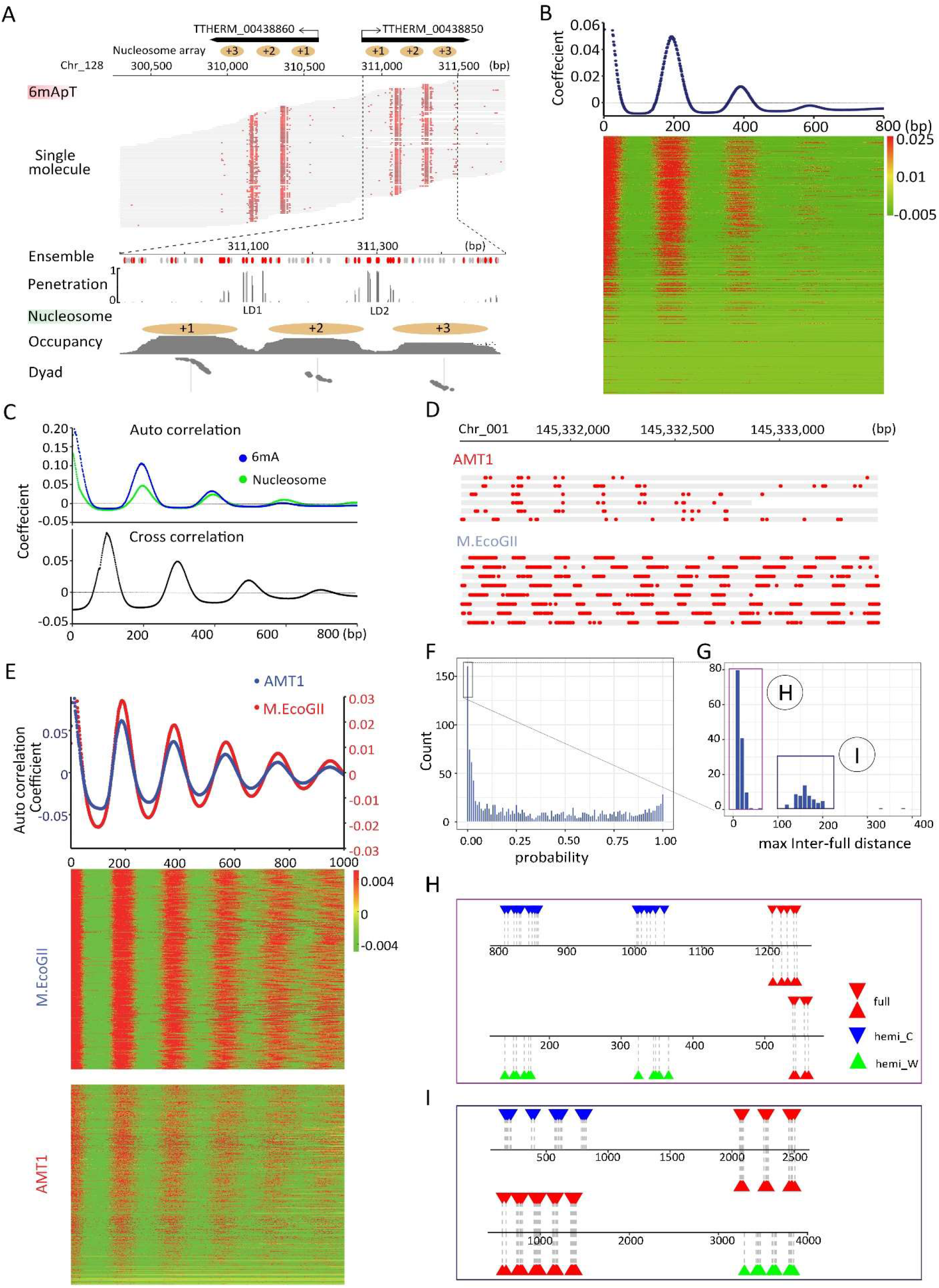
Chromatin-guided 6mA transmission. A. 6mA and nucleosome distributions in *Tetrahymena*. A typical genomic regions is shown with SMRT CCS reads mapped across it, as well as annotations of genes and canonical nucleosome arrays. Note that 6mApT sites (in either full or hemi-methylation, red dot) distributed along individual DNA molecules (gray line) are clustered in linker DNA (LD). LD1 is between the +1 and +2 nucleosome (the first and second nucleosome downstream of TSS); LD2 and beyond are defined iteratively further downstream of the gene body. B. Periodic 6mA distribution at the single molecule level in *Tetrahymena*. Autocorrelation between 6mA sites (distance≤1 kb) was calculated for individual DNA molecules, ranked by their median absolute deviations, and plotted as a heat map (bottom) and an aggregated correlogram (top). C. Autocorrelation of 6mA and nucleosome distributions at the ensemble level in *Tetrahymena* (top), revealing a ∼200bp periodicity. Cross-correlation between 6mA and nucleosome distributions (bottom), revealing a ∼100bp phase difference between them. D. Typical DNA molecules from human chromatin, after *in vitro* methylation by AMT1 complex and M.EcoGII, respectively. Note clusters of 6mA sites (red dot) distributed at regular intervals along individual DNA molecules (gray line). E. Periodic 6mA distributions at the single molecule level, after *in vitro* methylation by AMT1 complex and M.EcoGII, respectively. Autocorrelation between 6mA sites (distance ≤1kb) was calculated for individual DNA molecules, ranked by their median absolute deviations, and plotted as heat maps (bottom) and aggregated correlograms (top). F. Congregation of full-6mApT in DNA molecules undergoing hemi-to-full conversion. Their max inter-full distances were often very small, thus rarely represented (probability≤0.01) in simulations with permutated full and hemi positions (box); x-axis: the probability for simulated max inter-full distances to be no greater than the observed value; y-axis: the count of DNA molecules with the corresponding probability. G. Distribution of max inter-full distances for DNA molecules with strong full-6mApT congregation (probability≤0.01, Figure 5F box). Note the two peaks corresponding to DNA molecules with full-6mApT congregation within a LD (Figure 5H) and across adjacent LDs (Figure 5I), respectively. H. Full-6mApT congregation within a LD. I. Full-6mApT congregation across adjacent LDs.

We also analyzed human chromatin *in vitro* methylated by AMT1 complex or M.EcoGII (Figure 4D, E, Figure S7B-D). We first digested the DNA samples with *DpnI* (Figure S7A, B), targeting GATC sites with 6mA [43]. Only a fraction of GACT sites were cleaved, generating a nucleosome ladder strongly suggestive of preferential DNA methylation at linker DNA (Figure S7B). SMRT CCS revealed regularly distributed 6mA clusters on individual DNA molecules from both AMT1 complex and M.EcoGII-treated samples (Figure 5D). Autocorrelation analysis confirmed that 6mA sites were strongly phased, with cycles ranging from 160 to 200bp (Figure 5E: bottom). While 6mA density was substantially lower in the sample treated by AMT1 complex due to its strong preference for ApT sites (Figure 5D), the aggregated 6mA distribution correlogram showed the same cycle of ∼190bp for both samples (Figure 5E: top), underpinned by the nucleosome distribution pattern in human chromatin.

In contrast to *Tetrahymena* MAC genomic DNA, 6mA clusters on different DNA molecules from *in vitro* methylated human chromatin were poorly aligned for most genomic regions (Figure 5D). Indeed, 6mA distribution autocorrelation was much weaker for *in vitro* methylated human chromatin at the ensemble level (Figure S8B). In parallel, autocorrelation for nucleosome distribution at the ensemble level was much weaker in human than in *Tetrahymena*, indicating poor nucleosome positioning overall in human relative to *Tetrahymena* (Figure 4C: top, Figure S8B). As an exception that proves the rule, we found that around genomic positions with strong CTCF-binding, which are usually flanked by well-positioned nucleosomes [44, 45], 6mA sites from *in vitro* methylated human chromatin were strongly aligned, and importantly, 6mA peaks coincided with nucleosome troughs (Figure S8C). We conclude that mutual exclusivity between 6mA and the nucleosome is generally applicable at the single molecule level, but only manifests at the ensemble level for genomic regions with well-positioned nucleosomes. Our *in vitro* methylation results also indicate that preferential methylation of linker DNA is an intrinsic property for AMT1 complex, M.EcoGII, and potentially many other MTases.

### Processivity of AMT1-dependent methylation

Canonical maintenance MTases (*e*.*g*., *E. coli* dam DNA MTase) are generally processive rather than distributive [46]. In other words, upon substrate binding, they tend to catalyze multiple local methylation events before dissociation. To investigate processivity of AMT1-dependent methylation, we examined DNA molecules undergoing hemi-to-full conversion in WT *Tetrahymena* cells. We found that hemi-6mApT and full-6mApT distributions were often not random in these molecules (Figure 5F-I). Many exhibited full-6mApT congregation: the maximum observed distance between adjacent full-6mApT duplex positions (max inter-full distances) was much smaller than expected, and as a result rarely appeared in simulated controls, in which full-6mApT and hemi-6mApT positions were randomly permutated (Figure 5F). There was a strong tendency for multiple maintenance methylation events to occur in nearby positions. This tendency was especially prominent for DNA molecules early in the hemi-to-full conversion process, which were more likely to be methylated in a single processive run (Figure S9A).

For DNA molecules with strong full-6mApT congregation, their max inter-full distances were predominantly distributed in two peaks (Figure 5G): the left peak (max inter-full distances ≤30bp) corresponds to full-6mApT congregation within the same linker DNA (Figure 5H), while the right peak (130bp≤ max inter-full distances ≤200bp) corresponds to congregation across adjacent linker DNA regions (Figure 5I). In some DNA molecules, hemi-to-full conversion was already complete for one linker DNA (or at a higher level, gene), but not even started for its adjacent linker DNA region (or gene) (Figure 5H, I). More often, full-6mApT were intermixed with hemi-6mApT in one linker DNA region (or gene), while its adjacent linker DNA region (or gene) contained only hemi-6mApT (Figure S9B). The processivity of AMT1-dependent maintenance methylation therefore manifests as episodes of hemi-too-full conversion events that occur within one linker DNA (or gene), punctuated by switching of the MTase activity to its adjacent linker DNA (or gene).

### AMT1-independent *de novo* methylation

6mA levels were dramatically reduced but not eliminated in Δ*AMT1* cells (Table 1). Many ApT positions in the MAC genome were methylated in WT cells but not in Δ*AMT1* cells (Figure S10A). For genomic positions methylated in both, methylation penetration was generally much lower in Δ*AMT1* cells (Figure S10B). High penetration genomic positions were especially depleted in Δ*AMT1* cells (Figure 6A). Assuming exponential decay kinetics, we estimated the apparent half-life values for AMT1-dependent maintenance methylation (0.18× cell cycle) and AMT1-independent *de novo* methylation (4.2×) (Figure S10C). The fast AMT1-dependent maintenance methylation allows effective restoration of full-6mApT within one cell cycle in WT cells, while the slow AMT1-independent *de novo* methylation entails that in Δ*AMT1* cells, methylation plateau is only reached after multiple cell cycles. Indeed, in many DNA molecules from Δ*AMT1* cells, 6mA counts on W and C were disparate (Figure 2D: bottom, Figure 6B, Figure S10D). The strand with significantly fewer 6mA than expected for random distribution probably corresponds to the newly synthesized strand, which only carries 6mA newly deposited during the last cell cycle; the strand with significantly more 6mA probably corresponds to the old strand, which has accumulated 6mA over multiple cell cycles. The difficulty to propagate 6mA across the cell cycle also led to epigenetic instability in Δ*AMT1* cells, as different DNA molecules covering the same genomic region exhibited much higher variability of 6mA counts therein (Figure 6C).

**Figure 6.**
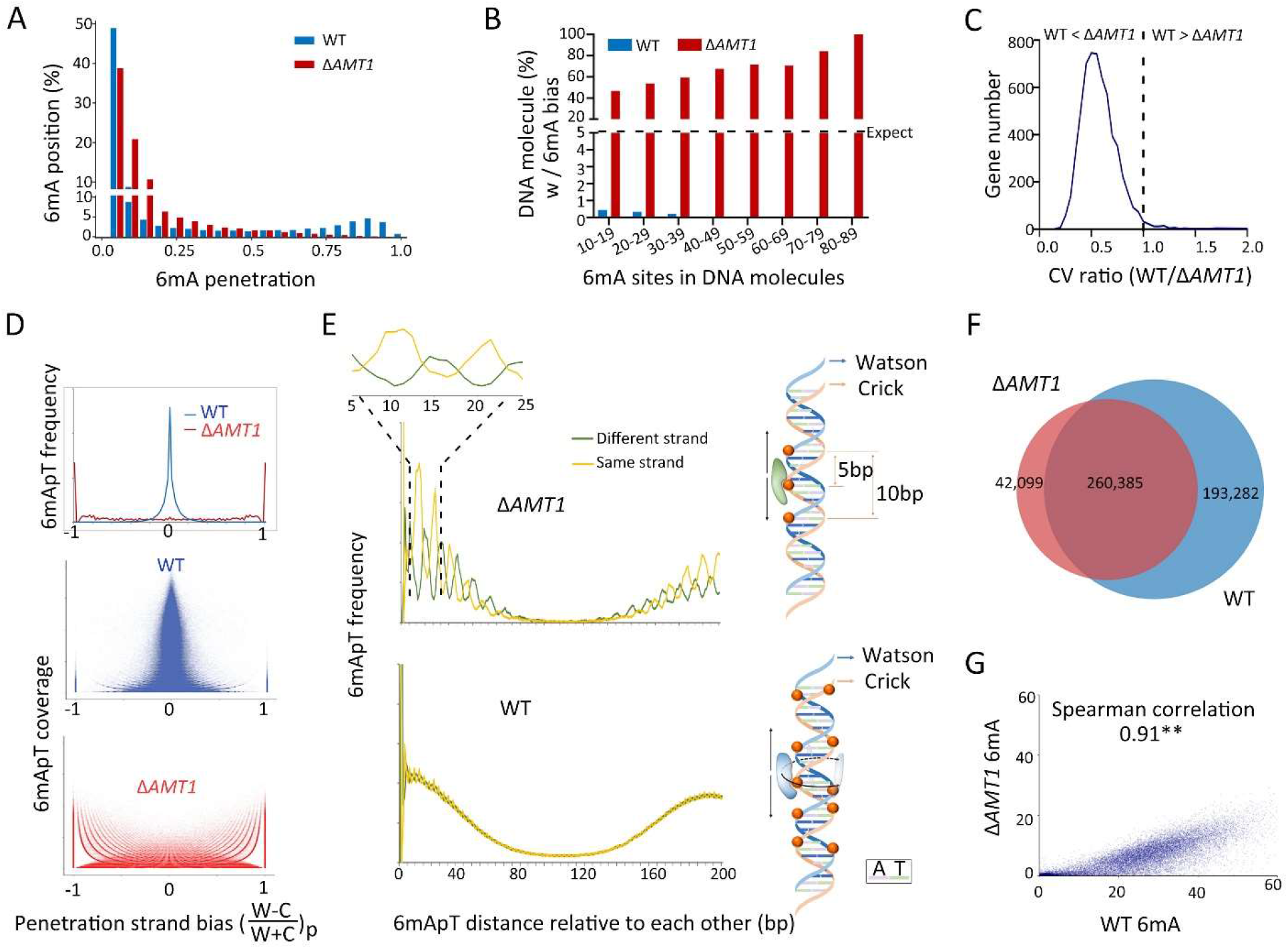
AMT1-independent *de novo* methylation. A. Depletion of high penetration 6mA positions in Δ*AMT1* relative to WT cells. B. Strong 6mA segregation strand biases in Δ*AMT1* cells. Chi-squared analysis was performed on DNA molecules with the specified number of total 6mA (full-6mApT counted as two, hemi-6mApT counted as one; x-axis), the percentage of DNA molecules with strong bias for 6mA segregation to one strand was indicated (expectance<5%, assuming random distribution; y-axis). WT cells were also analyzed as a negative control. C. Increased 6mA variability at the gene level in Δ*AMT1* relative to WT cells. For each gene, we calculated the coefficients of variance (CV) of 6mA counts from individual DNA molecules fully covering the gene, for WT and Δ*AMT1* cells, respectively. We then plotted the distribution of the ratio between the two CV Values 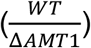 across all genes. Note that for most genes, the ratio is less than 1 (i.e., 6mA variability is higher in Δ*AMT1* than WT cells). D. Penetration strand bias of 6mA in WT and Δ*AMT1* cells. 6mA penetration strand bias is defined for an ApT position in the genome as the difference-sum ratio between the number of DNA molecules supporting 6mA on W and C, respectively: 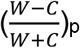. We plotted the distribution of ApT genomic positions according to their penetration strand bias (top). We also plotted their distribution according to both penetration strand bias and 6mA coverage (middle: WT; bottom: Δ*AMT1*). In WT cells, most ApT positions had penetration strand bias values around 0 (i.e., similar numbers of 6mA on W and C), while few had values at +1 (6mA only on W) or -1 (6mA only on C). The opposite was true for Δ*AMT1* cells. E. 10-bp cycle of 6mA penetration strand bias in Δ*AMT1* cells (top left), suggesting that the dedicated *de novo* 6mA-MTase can only approach the DNA substrate from one side (top right). Lack of such pattern in WT cells (bottom left) supports that AMT1 complex can approach from different sides (bottom right). F. Overlap in ApT positions methylated in WT or Δ*AMT1* cells (6mA penetration≥0.1). G. 6mA levels of individual genes in WT and Δ*AMT1* cells are strongly correlated. Each gene is assigned a coordinate: sum of 6mA penetration values for all methylated ApT positions in the gene body (ΣP) for WT (x-axis) and Δ*AMT1* cells (y-axis). The Spearman’s rank correlation coefficient is significant (p<0.01**).

In Δ*AMT1* cells, 6mA was also enriched in linker DNA and towards the 5’ end of Pol II-transcribed genes (Figure S11A, B). 6mApT sites, though present more sparsely, still formed clusters at regular intervals on individual DNA molecules (Figure 2D: bottom, Figure S11B). Autocorrelation analysis at both the single molecule level and the ensemble level showed a slight right shift in 6mA peaks, supporting increased linker DNA length (Figure S11C-E). We found many genomic regions that were more dispersively covered with 6mA in Δ*AMT1* than WT cells (Figure S11B). This may reflect reduced nucleosome positioning or increased nucleosome dynamics. In support, 6mA can directly promote nucleosome positioning, as the heavily methylated DNA becomes less bendable and thus prefers to be linker DNA rather than nucleosomal DNA [8, 10, 12]. Nucleosome positioning was indeed weakened in Δ*AMT1* relative to WT cells (Figure S11E) [11]. Alternatively, 6mA dispersion in Δ*AMT1* cells may be attributed to the slow AMT1-independent *de novo* methylation, which records nucleosome movement throughout the cell cycle rather than only briefly after DNA replication.

In strong contrast to WT cells, 6mA penetration for most ApT positions in the MAC genome of Δ*AMT1* cells showed strong biases for either W or C, and many were exclusively methylated on one strand (Figure 2D: bottom, Figure 6D); this tendency grew in prominence with increasing sequencing coverage (Figure 6D: bottom), thus unlikely an artefact of random fluctuations. Intriguingly, genomic positions with strong penetration bias for W or C exhibited periodic distributions with a ∼10-bp cycle (Figure 6E). This matches the pitch of the DNA double helix, suggesting that the dedicated *de novo* MTase is constrained to approach the DNA substrate from only one side (Figure 6E). The strong penetration bias also precludes this MTase from playing a major role in maintenance methylation.

Despite these distinctions, there were also connections between AMT1-dependent and AMT1-independent methylation. Most ApT positions methylated in Δ*AMT1* cells were also methylated in WT cells; the two sets essentially converged at high methylation penetration (Figure 6F, Figure S10A). Furthermore, 6mA levels at individual genes and even individual linker DNA regions of a gene showed strong correlations between WT and Δ*AMT1* cells (Figure 6G and S10E). These connections suggest an integrated 6mA transmission pathway: AMT1-independent *de novo* methylation primes the system by laying down an incipient 6mApT distribution pattern, which is fulfilled and transmitted by AMT1-dependent maintenance methylation.

## Discussion

### 6mA detection by SMRT CCS

In the free 6mA nucleotide, N^6^-methyl group minimizes the steric clash by pointing towards the Watson-Crick edge of the purine ring [6]. This is likely also the preferred conformation in single-stranded DNA. However, in double-stranded DNA, N^6^-methyl group must adopt the energetically less favorable conformation and point the other way, to allow the N^6^-lone pair electrons to engage in Watson-Crick base pairing. This entails a pause in DNA synthesis, as the DNA polymerase waits for 6mA in the template strand to switch conformation. In SMRT sequencing, this is recorded as increased IPD. SMRT CCS allows robust evaluation of IPD at the single site and single molecule level, as multiple passes by the DNA polymerase overcome random fluctuations. We rationalize that 6mA and unmodified A feature distinct IPDr distributions, which can be deconvoluted effectively. Based on these basic assumptions, we have developed a bioinformatic pipeline to fully exploit the recent progress in SMRT CCS for strand-specific, accurate, and sensitive detection of 6mA on individual DNA molecules multi-kb in length.

6mA detection by SMRT CCS is critical for our study in the following aspects. First, SMRT CCS detects 6mA with high accuracy (low false positive rates), which allows us to ***1)*** determine the ApT specificity for AMT1-dependent maintenance methylation and AMT1-independent *de novo* methylation, and ***2)*** distinguish hemi-6mApT from full-6mApT in WT and Δ*AMT1* cells. Second, SMRT CCS preserves long-range connectivity information at the single molecule level, which allows us to ***1)*** identify hemi^+^/BrdU^+^ molecules and establish segregation of hemi-6mApT to the old strand after DNA replication, and ***2)*** identify DNA molecules undergoing maintenance methylation and characterize AMT1 processivity. Third, SMRT CCS detects 6mA with high sensitivity (low false negative rates), which, combined with deep sequencing coverage of the *Tetrahymena* MAC genome, allows us to ***1)*** unambiguously identify rare methylation events, and ***2)*** to generate absolute and exact quantification of 6mA levels over a genomic region. Furthermore, there is gross discrepancy between CLR and CCS-based assessments of many key 6mA parameters in *Tetrahymena* cells, including percentage of 6mA in non-ApT context, percentage of hemi- and full-6mApT, and percentage of ApT positions with 6mA penetration bias. In all cases, the misleading CLR results are likely rooted in erroneous extrapolation from the ensemble level. Without SMRT CCS, most of our conclusions simply cannot be drawn.

As a gold standard for 6mA detection, SMRT CCS boast some outstanding features: *1)* low background noise (∼0.01%), ***2)*** high accuracy (∼99%), ***3)*** high sensitivity (∼99%), and ***4)*** long read (∼3kb). It is worth noting that these parameters are mutually connected and can be individually optimized according to one’s need. At the cost of CCS read length/DNA insert size (and consequently, sequencing coverage), we chose to increase the number of CCS passes to improve the first three parameters. Shifting IPDr threshold for 6mA calling affects accuracy and sensitivity of 6mA detection in the opposite direction. Our deconvolution-based approach automatically set the threshold to achieve a balanced outcome. SMRT CCS can be used to detect other base modifications, such as BrdU. While we limited ourselves to a single readout of the DNA polymerase kinetics (IPD) and a rationally designed algorithm (independent of ground truth training data), there is great potential in incorporating additional readout and implementing neural network-based machine learning algorithms [47].

6mA is highly enriched in linker DNA in *Tetrahymena*. The resulting 6mA clusters, regularly spaced, demarcate individual nucleosomes on a chromatin fiber, providing long-range epigenetic information generally missing from short-read sequencing data. Specific methylation of linker DNA is likely an intrinsic feature of AMT1 complex, M.EcoGII, and many other 6mA-MTases. This property can be exploited to probe chromatin organization via *in vitro* methylation, analogous to nuclease protection [48-51]. This is an especially powerful approach when combined with 6mA detection by SMRT CCS [48, 50].

### AMT1-dependent methylation

We have extensively characterized AMT1-dependent 6mA transmission. Our *in vivo* results demonstrate high specificity for maintenance methylation at the ApT dinucleotide, while our *in vitro* results support substantial *de novo* methylation activity at ApT sites and, to a lesser degree, non-ApT sites. Note that 6mA at non-ApT sites is necessarily the product of *de novo* methylation. We emphasize that *de novo* methylation underpins the biochemical assay by which the *Tetrahymena* MTase activity and eventually AMT1 complex were identified [10, 32]. Indeed, DNMT1, the eukaryotic maintenance MTase required for semi-conservative transmission of 5-methylcytosine (5mC) in the CpG dinucleotide, also has *de novo* methylation activity [52, 53]. We argue that AMT1-dependent *de novo* methylation is amplified under *in vitro* conditions, while curtailed by various *in vivo* circumstances. ***1)*** For *in vitro* methylation of human chromatin, *de novo* methylation precedes—and is the prerequisite for—maintenance methylation. In contrast, the abundance of hemi-6mApT in *Tetrahymena* MAC DNA immediately after DNA replication allows maintenance methylation to effectively outcompete *de novo* methylation *in vivo*. ***2)*** Processivity of AMT1-dependent methylation may enhance the preference for maintenance methylation *in vivo*. Multiple hemi-6mApT sites, present in a cluster often fully covering a linker DNA, are readily converted to full-6mApT with little chance of *de novo* methylation as the side reaction. ***3)*** AMT1-dependent maintenance methylation may be further enhanced by other *in vivo* factors. In *Tetrahymena*, 6mA is highly enriched in linker DNA flanked by nucleosomes containing H3K4me3 and H2A.Z [7], which may interact with AMT1 complex and modulate its substrate specificity.

### Comparing 6mA and 5mC pathways in eukaryotes

Our work provides definitive evidence for a eukaryotic 6mA pathway comprising two distinct but linked steps: AMT1-independent *de novo* methylation and AMT1-dependent maintenance methylation (Figure 7). While AMT1-independent *de novo* methylation is dispensable for maintaining the 6mA pattern in the MAC of asexually propagating *Tetrahymena* cells [11], it is likely to play a critical role during sexual reproduction, as the transcriptionally silent and 6mA-free germline MIC is differentiated into the transcriptionally active and 6mA-rich somatic MAC. This two-step pathway bears some striking resemblance to the eukaryotic 5mC pathway, featuring the DNMT3A/3B-dependent *de novo* methylation and DNMT1-dependent maintenance methylation for transmission of 5mC at the CpG dinucleotide (Figure 7) [54]. As *bona fide* eukaryotic epigenetic marks, 6mA and 5mC play opposite roles in transcription regulation (Figure 7). Their transmission pathways are deep-rooted and widespread, but show distinct phylogenetic distributions, with homologues of AMT1 complex components notably missing from land plants, higher fungi, and animals (Figure S12). 6mA and 5mC therefore represent a pair of critical switches that can dramatically alter the global transcription landscape, and their presence or loss may drive some major branching events in eukaryotic evolution.

**Figure 7.**
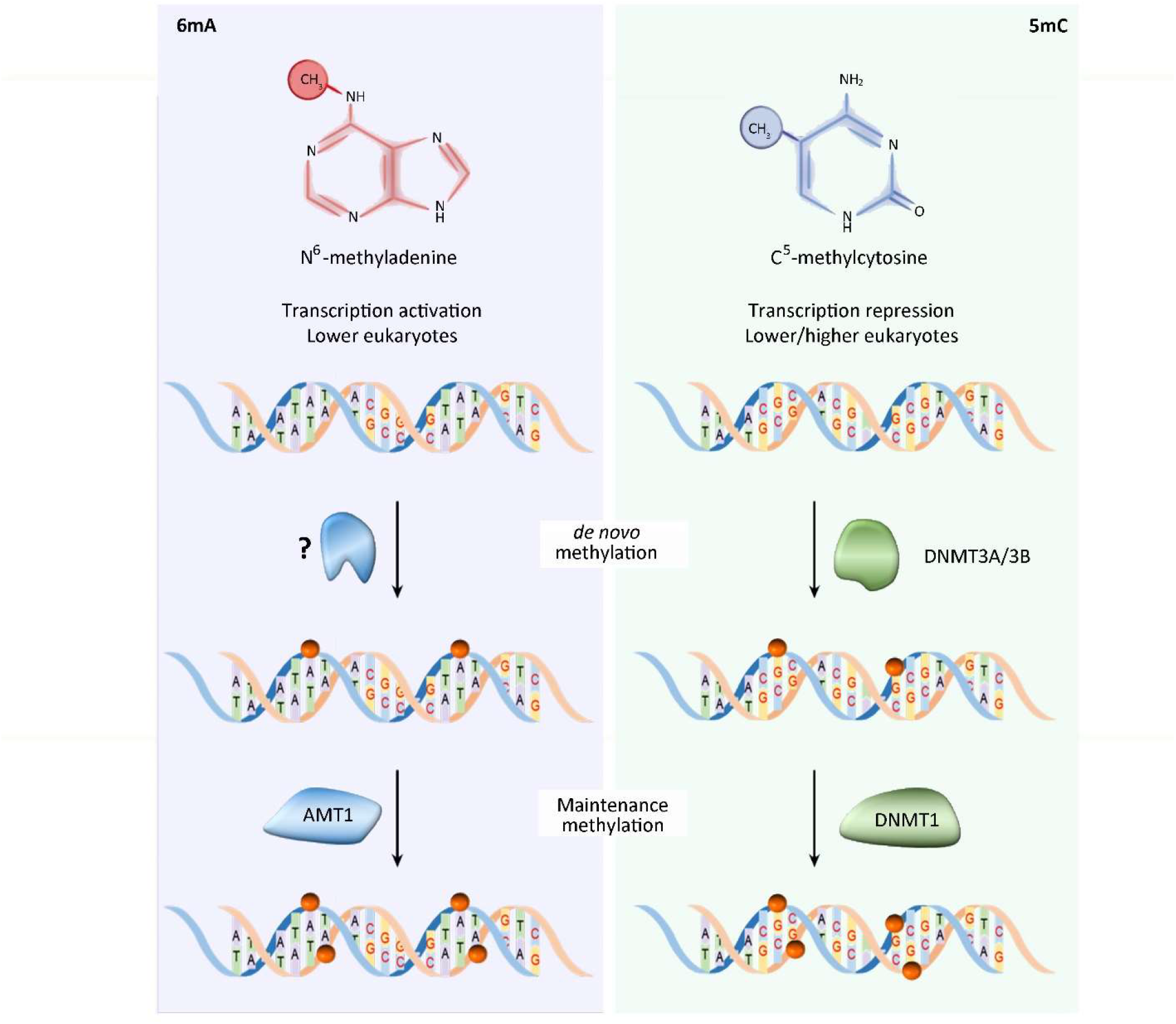
Comparison of 6mA and 5mC pathways in eukaryotes. See text for details.

## Conclusions

A SMRT CCS-based pipeline is developed for strand-specific, accurate, and sensitive detection of 6mA on individual DNA molecules multi-kb in length. It is implemented to achieve genome-wide mapping of 6mA distribution at the single molecule level in *Tetrahymena*, the first eukaryotic system to be so thoroughly characterized. Combined with SMRT CCS-based BrdU detection, this provides a rigorous proof that 6mA is transmitted by a semi-conservative mechanism: full-6mApT is split by DNA replication into hemi-6mApT segregated to the old strand, which is restored to full-6mApT by AMT1-dependent maintenance methylation. This mechanism is probably conserved in unicellular eukaryotes like protists, green algae, and basal fungi, making 6mA a *bona fide* epigenetic mark in these systems. Furthermore, dissection of AMT1-dependent maintenance methylation and AMT1-independent *de novo* methylation reveals a molecular pathway for 6mA transmission with striking similarity to 5-methyl cytosine (5mC) transmission at the CpG dinucleotide, with implications in how epigenetic regulation of global transcription may shape and be shaped by eukaryotic evolution.

## Materials and Methods

Additional details are available in Supplemental Material.

### *Tetrahymena* strains

*Tetrahymena thermophila* WT strain (SB210) was obtained from the *Tetrahymena* Stock Center. Δ*AMT1* was a homozygous homokaryon (MAC and MIC) knockout strain generated in our previous study [11].

### *In vitro* reconstitution of AMT1 complex

The DNA sequences encoding the *Tetrahymena* AMT1, AMT7, AMTP1 (1-240 aa, truncating the C-terminal low complexity region that may interfere with overexpression and purification) and AMTP2 proteins were each codon optimized for *E. coli* expression and synthesized. AMT1 and AMT7 were inserted in tandem into an in-house bacterial expression vector, in which a His6-MBP tag was fused to the AMT1 sequence via a TEV protease cleavage site. AMTP1 (1-240) and AMTP2 were cloned to a modified pRSF-Duet vector for co-expression, with AMTP2 preceded by an N-terminal His_6_-SUMO tag and a ubiquitin-like protease 1 (ULP1) cleavage site. BL21(DE3) RIL cells harboring the expression plasmids were grown at 37°C and induced by addition of 0.2 mM isopropyl β-D-1-thiogalactopyranoside (IPTG) when the cell density reached A_600_ of 1.0. The cells continued to grow at 16°C overnight. Subsequently, the cells were harvested and lysed in buffer containing 50mM Tris-HCl (pH 8.0), 1M NaCl, 25mM Imidazole, 10% glycerol and 1mM PMSF. The fusion proteins were purified through nickel affinity chromatography, followed by removal of His_6_-MBP and His_6_-SUMO tags by TEV and ULP1 cleavage, ion-exchange chromatography on a Heparin column (GE Healthcare), and size-exclusion chromatography on a 16/600 Superdex 200pg column (GE Healthcare). The purified proteins were concentrated in 20mM Tris-HCl (pH 7.5), 100mM NaCl, 5% glycerol and 5mM DTT, and stored at -80°C.

### DNA methyltransferase assay of reconstituted AMT1 complex

Synthesized 12-mer DNA oligos containing one central 6mA-modified ApT site were annealed to generate the substrates (upper strand: 5′-GCA AG(6mA) TCA ACG -3′, lower stand: 5′-CGT TGA TCT TGC -3′). For steady-state kinetic assay, a 20μL reaction mixture contained the hemi-methylated substrate at various concentrations (0, 0.04, 0.1, 0.16, 0.24, 0.36, 0.5, 0.75, 1, 1.5, 2μM), 0.01μM AMT1 complex, 0.55μM S-adenosyl-L-[methyl-^3^H] methionine (specific activity 18Ci/mmol, PerkinElmer), 1.9μM AdoMet in 50mM Tris–HCl, pH 8.0, 0.05% β-mercaptoethanol, 5% glycerol, and 200μg/mL BSA. For substrate specificity assay, a 15μL reaction mixture contained 2μM unmodified or hemi-methylated substrates, 0.1μM AMT1 complex, 3μM S-adenosyl-L-[methyl-^3^H] methionine (specific activity 18Ci/mmol, PerkinElmer). For unmodified DNA duplex, upper strand: 5′-AAC TTC TGT CAT TAC ATT AAG CTT TAA -3′, lower stand: 5′-TTA AAG CTT AAT GTA ATG ACA GAA GTT -3′. For hemi-methylated DNA duplex, upper strand: 5′-AAC TTC TGT C(6mA)T TAC (6mA)TT AAG CTT TAA -3′, lower stand: 5′-TTA AAG CTT AAT GTA ATG ACA GAA GTT -3′. The assays were performed in triplicate at room temperature for 30 min.

### *In vitro* methylation of human chromatin

×10^5^ OCI-AML3 cells were lysed in 0.5 mL of nuclei extraction buffer (20mM HEPES pH 7.9, 10mM KCl, 0.1% Triton X-100, 20% glycerol, 0.5mM spermidine, 1× Protease Inhibitor Cocktail) for 8 min on ice. Purified nuclei were methylated in a 30μL of 50mM Tris-HCl pH 8.0, 2mM EDTA, 0.5mM EDGA, 160μM SAM, 1× Protease Inhibitor Cocktail and 38μM AMT1-complex or M.EcoGII or no enzyme control for 1h at 37°C. Genomic DNA was extracted with Monarch® HMW DNA Extraction Kit for Cells & Blood (NEB), digested with DpnI overnight at 37°C, and resolved on 1% agarose gel. DNA fragments 3-5kb in length were gel purified for SMRT sequencing.

### SMRT sequencing sample preparation

Smaller insert size favors accurate calling of 6mA as well as regular bases at the single molecule level, which is dependent on the number of CCS passes of individual DNA molecules. Larger insert size (generating more sequenced bp per SMRT Cell given the fixed upper limit for total reads) favors sensitive calling of low penetration 6mA positions at the ensemble level, which is dependent on the overall sequencing coverage of the target genome. As a compromise, we chose genomic DNA fragments 3-5kb in length (generated by either sonication or *Dpn*I digestion) for SMRT CCS library preparation, so that most DNA molecules had ≥30 CCS passes, given the read length distribution on the Sequel II platform (N50≥100kb).

Genomic DNA was extracted from *Tetrahymena* WT (SB210, with or without BrdU-labeling) and Δ*AMT1* cells using Wizard® Genomic DNA Purification Kit (Promega, A1120), sheared to 3-5kb in length with Megaruptor (Diagenode Diagnostics), and used to generate sequencing libraries for the Sequel II System. For BrdU-labeling, G1-synchronized *Tetrahymena* cells were collected by centrifugal elutriation [55], released into the fresh medium with 0.4mM BrdU, and collected after 0h, 1.5h, 2h, or 4h for genomic DNA extraction and SMRT sequencing.

### SMRT CCS data analysis

Single molecule sam files were extracted from the SMRT sequencing data using custom Perl script and transformed into single molecule bam files by samtools [56]. Circular Consensus Sequence (CCS) was calculated for each DNA molecule using the CCS module (SMRT Link v10.2, Pacific Biosciences). Only DNA molecules with high subread coverage (≥30×) were retained. Single molecule aligned bam files were generated using BLASR [57], which in turn served as the input for the ipdSummary module to calculate IPD ratios (IPDr). Self-referencing not only allows 6mA calling at the single molecule level, but also greatly speeds up computation.

We removed DNA molecules with global dispersion of IPDr for unmodified adenine sites: IPDr standard deviation (SD) ≥0.35. We also removed DNA molecules with local dispersion of IPDr, referred to as N* clusters. N= G, C, T; N*: IPDr≥2.8; N* cluster: inter-N* distances ≤25bp, N* count ≥4, on the same strand. Finally, an IPDr threshold was determined by peak deconvolution (IPDr=2.38 for Replicate 1 of WT cells) for calling bulk 6mA in the remaining DNA molecules.

For calling BrdU, we adopted a similar pipeline with the following modifications. Regions adjacent to 6mApT sites (both strands: -10 to +10bp) were masked from further analysis to avoid interference between 6mA and BrdU. IPDr 2.5 was set as the threshold for calling BrdU. Note that BrdU^+^ molecules only represent a small fraction of BrdU-labeled DNA molecules (∼10% of SMRT CCS reads were BrdU^+^ in synchronized G2 cells with near complete BrdU-labeling), due to the high threshold (BrdU sites ≥15) and the high false negative rate of BrdU calls.

CCS reads were mapped back to the *Tetrahymena* reference sequences for the MAC, MIC, and mitochondrion (merged into a single file) by blastn [58]. The parameters “-max-target_seqs 2, -max_hsps 2” were used to allow identification and mapping of bipartite reads (with two segment pairs). We focused on fully mapped reads (mapped length ≥98% for the only segment pair or segment pairs combined). Segment pairs with mapped identity ≥95% were retained for further analysis. DNA molecules mapped specifically to the MIC rather than the MAC—usually containing MIC-limited internal eliminated sequences (IES) comprising transposable elements and repetitive sequences—were distinguished by much higher blastn alignment scores (Δ≥50) matching the MIC reference genome than the MAC. Some DNA molecules fully mapped to the MAC genome may come from the MIC, as they correspond to genomic regions not interrupted by MIC-limited sequences. Nonetheless, due to the high ploidy (∼90×) of the MAC relative to the diploid MIC [59], they only represent a small fraction (<5%), thus should not significantly affect the analysis results.

### Penetration strand bias and segregation strand bias

6mA penetration strand bias, 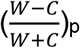, is defined for an ApT position in the genome as the difference-sum ratio between the number of DNA molecules supporting 6mA on W and C, respectively (W+C≥10). The values range between -1 (6mA only on C) and 1 (6mA only on W). 6mA penetration strand bias was calculated for both WT and Δ*AMT1* cells. We identified ApT positions with penetration strand bias of +1 or -1 in Δ*AMT1* cells and generated phasogram (defined as histogram of distances between specified positions) to reveal their periodic distribution relative to each other.

6mA segregation strand bias, 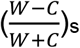, is defined for a hemi^+^ molecule (W+C≥11 or W||C≥11) as the difference-sum ratio between the count of hemi-6mApT on W and C, respectively. The values range between -1 (only hemi-C) and 1 (only hemi-W). Note that 6mA in full-6mApT is not included in this calculation. Segregation strand bias is also calculated for BrdU sites in BrdU^+^ molecules (BrdU≥15 or W||C≥15).

### Autocorrelation and cross-correlation analysis

A vector consisting of a series of 0’s (no 6mA) and 1’s (6mA at the position on either strand) was encoded for each DNA molecule with 6mA sites (≥2). This vector was the input for the acf function in the statsmodel python package [60] for computing 6mA autocorrelation coefficients at the single molecule level. For correlation analysis at ensemble level, the MAC reference genome was divided into 5kb regions. We then used bedtools coverage subcommand [61] to count 6mA or nucleosome dyad across all genomic positions, generating two encoding vectors for each such genomic region (focusing on those with 6mA genomic positions: ≥2; 6mA genomic positions supported by only one 6mA site/DNA molecule are excluded to reduce background). This pair of vectors were the input for the acf function for computing the autocorrelation coefficients of 6mA and nucleosome distributions, respectively; they were also the input for the ccf function for computing the cross-correlation coefficients between 6mA and nucleosome distributions.

### Full-6mApT congregation

For each DNA molecule undergoing hemi-to-full conversion (full-6mApT ≥5, hemi-6mApT ≥9), we first calculated the observed maximum value of distances between adjacent full-6mApT duplexes (*D*_*obs*_). We then calculated the equivalent values for 1000 simulations, in which the full-6mApT and hemi-6mApT positions in the same DNA molecule were randomly permutated (*D*_*sim*_). This allowed us to estimate the probability for observed full-6mApT congregation (*D*_*sim*_ ≤ *D*_*obs*_), assuming that maintenance methylation is random.

## Acknowledgment

*Tetrahymena* WT strains were obtained from the *Tetrahymena* Stock Center (http://tetrahymena.vet.cornell.edu). SMRT sequencing was performed at the Genomics Core Facility, Icahn School of Medicine at Mount Sinai. High-performance computing resources for data processing were provided by the Institute of Evolution and Marine Biodiversity at Ocean University of China, the Center for High Performance Computing and System Simulation at Laoshan Laboratory, and the Center for Advanced Research Computing (CARC) at the University of Southern California.

## Funding

National Natural Science Foundation of China 32200336 (Y.S.), National Natural Science Foundation of China 32200399 (Y.W.), National Natural Science Foundation of China 32125006, 32070437 (S.G.), Department of Biochemistry and Molecular Medicine at University of Southern California (Y.L.).

## Author contributions

Yifan L. and S.G. conceived the project, supervised most of the experiments and data analyses, and prepared the manuscript. Y.S. and W.Y. analyzed SMRT CCS data. Y.W. and Yongqiang L. performed the *Tetrahymena* experiments. J.L. (supervised by J.S.) and B.N. (supervised by S.G.) reconstituted and characterized AMT1 complex, which was used by X.Q.W. (supervised by Y.D.) to perform *in vitro* methylation of the human chromatin. B.P. analyzed Illumina sequencing data and phylogenetic distribution of DNA MTases. C.L. performed statistical analyses.

### Competing interests

None.

## Notes

### Competing Interest Statement

The authors have declared no competing interest.

## References

1. Shi H, Wei J, He C. Where, When, and How: Context-Dependent Functions of RNA Methylation Writers, Readers, and Erasers. Mol Cell. 2019;74(4):640–650.

2. Ignatova VV, Stolz P, Kaiser S, Gustafsson TH, Lastres PR, Sanz-Moreno A, et al. The rRNA m(6)A methyltransferase METTL5 is involved in pluripotency and developmental programs. Genes Dev. 2020;34(9-10):715–729.

3. Saneyoshi M, Harada F, Nishimura S. Isolation and characterization of N6-methyladenosine from Escherichia coli valine transfer RNA. Biochim Biophys Acta. 1969;190(2):264–273.

4. Iwanami Y, Brown GM. Methylated bases of ribosomal ribonucleic acid from HeLa cells. Arch Biochem Biophys. 1968;126(1):8–15.

5. Sanchez-Romero MA, Cota I, Casadesus J. DNA methylation in bacteria: from the methyl group to the methylome. Curr Opin Microbiol. 2015;25:9–16.

6. Bochtler M, Fernandes H. DNA adenine methylation in eukaryotes: Enzymatic mark or a form of DNA damage? Bioessays. 2021;43(3):e2000243.

7. Wang Y, Chen X, Sheng Y, Liu Y, Gao S. N6-adenine DNA methylation is associated with the linker DNA of H2A.Z-containing well-positioned nucleosomes in Pol II-transcribed genes in Tetrahymena. Nucleic Acids Res. 2017;45(20):11594–11606.

8. Fu Y, Luo GZ, Chen K, Deng X, Yu M, Han D, et al. N6-methyldeoxyadenosine marks active transcription start sites in Chlamydomonas. Cell. 2015;161(4):879–892.

9. Mondo SJ, Dannebaum RO, Kuo RC, Louie KB, Bewick AJ, LaButti K, et al. Widespread adenine N6-methylation of active genes in fungi. Nat Genet. 2017;49(6):964–968.

10. Beh LY, Debelouchina GT, Clay DM, Thompson RE, Lindblad KA, Hutton ER, et al. Identification of a DNA N6-adenine methyltransferase complex and its impact on chromatin organization. Cell. 2019;177(7):1781–1796.

11. Wang Y, Sheng Y, Liu Y, Zhang W, Cheng T, Duan L, et al. A distinct class of eukaryotic MT-A70 methyltransferases maintain symmetric DNA N6-adenine methylation at the ApT dinucleotides as an epigenetic mark associated with transcription. Nucleic Acids Res. 2019;47(22):11771–11789.

12. Luo GZ, Hao Z, Luo L, Shen M, Sparvoli D, Zheng Y, et al. N(6)-methyldeoxyadenosine directs nucleosome positioning in Tetrahymena DNA. Genome Biol. 2018;19(1):200.

13. Greer EL, Blanco MA, Gu L, Sendinc E, Liu J, Aristizabal-Corrales D, et al. DNA methylation on N6-adenine in C. elegans. Cell. 2015;161(4):868–878.

14. Koziol MJ, Bradshaw CR, Allen GE, Costa ASH, Frezza C, Gurdon JB. Identification of methylated deoxyadenosines in vertebrates reveals diversity in DNA modifications. Nat Struct Mol Biol. 2016;23(1):24–30.

15. Liang Z, Shen L, Cui X, Bao S, Geng Y, Yu G, et al. DNA N(6)-adenine methylation in Arabidopsis thaliana. Dev Cell. 2018;45(3):406–416 e403.

16. Liu J, Zhu Y, Luo GZ, Wang X, Yue Y, Wang X, et al. Abundant DNA 6mA methylation during early embryogenesis of zebrafish and pig. Nat Commun. 2016;7:13052.

17. Wang X, Li Z, Zhang Q, Li B, Lu C, Li W, et al. DNA methylation on N6-adenine in lepidopteran Bombyx mori. Biochim Biophys Acta Gene Regul Mech. 2018;1861(9):815–825.

18. Wu TP, Wang T, Seetin MG, Lai Y, Zhu S, Lin K, et al. DNA methylation on N(6)-adenine in mammalian embryonic stem cells. Nature. 2016;532(7599):329–333.

19. Xiao CL, Zhu S, He M, Chen Zhang Q, Chen Y, et al. N(6)-Methyladenine DNA Modification in the Human Genome. Mol Cell. 2018;71(2):306–318 e307.

20. Zhang G, Huang H, Liu D, Cheng Y, Liu X, Zhang W, et al. N6-methyladenine DNA modification in Drosophila. Cell. 2015;161(4):893–906.

21. Zhou C, Wang C, Liu H, Zhou Q, Liu Q, Guo Y, et al. Identification and analysis of adenine N(6)-methylation sites in the rice genome. Nat Plants. 2018;4(8):554–563.

22. Kong Y, Cao L, Deikus G, Fan Y, Mead EA, Lai W, et al. Critical assessment of DNA adenine methylation in eukaryotes using quantitative deconvolution. Science. 2022;375(6580):515–522.

23. Schiffers S, Ebert C, Rahimoff R, Kosmatchev O, Steinbacher J, Bohne AV, et al. Quantitative LC-MS provides no evidence for m(6)dA or m(4)dC in the genome of mouse embryonic stem cells and tissues. Angew Chem Int Ed Engl. 2017;56(37):11268–11271.

24. Musheev MU, Baumgartner A, Krebs L, Niehrs C. The origin of genomic N(6)-methyl-deoxyadenosine in mammalian cells. Nat Chem Biol. 2020;16(6):630–634.

25. Liu X, Lai W, Li Y, Chen S, Liu B, Zhang N, et al. N(6)-methyladenine is incorporated into mammalian genome by DNA polymerase. Cell Res. 2021;31(1):94–97.

26. Iyer LM, Zhang D, Aravind L. Adenine methylation in eukaryotes: Apprehending the complex evolutionary history and functional potential of an epigenetic modification. Bioessays. 2016;38(1):27–40.

27. Iyer LM, Abhiman S, Aravind L. Natural history of eukaryotic DNA methylation systems. Prog Mol Biol Transl Sci. 2011;101:25–104.

28. Liu J, Yue Y, Han D, Wang X, Fu Y, Zhang L, et al. A METTL3-METTL14 complex mediates mammalian nuclear RNA N6-adenosine methylation. Nat Chem Biol. 2014;10(2):93–95.

29. Gorovsky MA, Hattman S, Pleger GL. (6N)methyl adenine in the nuclear DNA of a eucaryote, Tetrahymena pyriformis. J Cell Biol. 1973;56(3):697–701.

30. Meyer E, Chalker DL. Epigenetics of ciliates. In Epigenetics. Edited by Allis CD, Jenuwein T, Reinberg D, Caparros M: Cold spring harbor laboratory press; 2007: 127–150

31. Karrer KM, VanNuland TA. Methylation of adenine in the nuclear DNA of Tetrahymena is internucleosomal and independent of histone H1. Nucleic Acids Res. 2002;30(6):1364–1370.

32. Bromberg S, Pratt K, Hattman S. Sequence specificity of DNA adenine methylase in the protozoan Tetrahymena thermophila. J Bacteriol. 1982;150(2):993–996.

33. Karrer KM. Nuclear dualism. Methods Cell Biol. 2012;109:29–52.

34. Xu J, Zhao X, Mao F, Basrur V, Ueberheide B, Chait BT, et al. A Polycomb repressive complex is required for RNAi-mediated heterochromatin formation and dynamic distribution of nuclear bodies. Nucleic Acids Res. 2021;49(10):5407–5425.

35. Zhao X, Xiong J, Mao F, Sheng Y, Chen X, Feng L, et al. RNAi-dependent Polycomb repression controls transposable elements in Tetrahymena. Genes Dev. 2019;33(5-6):348–364.

36. Flusberg BA, Webster DR, Lee JH, Travers KJ, Olivares EC, Clark TA, et al. Direct detection of DNA methylation during single-molecule, real-time sequencing. Nat Methods. 2010;7(6):461–465.

37. Eid J, Fehr A, Gray J, Luong K, Lyle J, Otto G, et al. Real-time DNA sequencing from single polymerase molecules. Science. 2009;323(5910):133–138.

38. Wenger AM, Peluso P, Rowell WJ, Chang PC, Hall RJ, Concepcion GT, et al. Accurate circular consensus long-read sequencing improves variant detection and assembly of a human genome. Nat Biotechnol. 2019;37(10):1155–1162.

39. Chen J, Hu R, Chen Y, Lin X, Xiang W, Chen H, et al. Structural basis for MTA1c-mediated DNA N6-adenine methylation. Nat Commun. 2022;13(1):3257.

40. Murray IA, Morgan RD, Luyten Y, Fomenkov A, Correa IR, Jr., Dai N, et al. The non-specific adenine DNA methyltransferase M.EcoGII. Nucleic Acids Res. 2018;46(2):840–848.

41. Douvlataniotis K, Bensberg M, Lentini A, Gylemo B, Nestor CE. No evidence for DNA N (6)-methyladenine in mammals. Sci Adv. 2020;6(12):eaay3335.

42. O’Brown ZK, Boulias K, Wang J, Wang SY, O’Brown NM, Hao Z, et al. Sources of artifact in measurements of 6mA and 4mC abundance in eukaryotic genomic DNA. BMC Genom. 2019;20(1):445.

43. Vovis GF, Lacks S. Complementary action of restriction enzymes endo R-DpnI and Endo R-DpnII on bacteriophage f1 DNA. J Mol Biol. 1977;115(3):525–538.

44. Fu Y, Sinha M, Peterson CL, Weng Z. The insulator binding protein CTCF positions 20 nucleosomes around its binding sites across the human genome. PLoS Genet. 2008;4(7):e1000138.

45. Krietenstein N, Abraham S, Venev SV, Abdennur N, Gibcus J, Hsieh TS, et al. Ultrastructural details of mammalian chromosome architecture. Mol Cell. 2020;78(3):554–565 e557.

46. Urig S, Gowher H, Hermann A, Beck C, Fatemi M, Humeny A, et al. The Escherichia coli dam DNA methyltransferase modifies DNA in a highly processive reaction. J Mol Biol. 2002;319(5):1085–1096.

47. Tse OYO, Jiang P, Cheng SH, Peng W, Shang H, Wong J, et al. Genome-wide detection of cytosine methylation by single molecule real-time sequencing. Proc Natl Acad Sci U S A. 2021;118(5).

48. Stergachis AB, Debo BM, Haugen E, Churchman LS, Stamatoyannopoulos JA. Single-molecule regulatory architectures captured by chromatin fiber sequencing. Science. 2020;368(6498):1449–1454.

49. Altemose N, Maslan A, Smith OK, Sundararajan K, Brown RR, Mishra R, et al. DiMeLo-seq: a long-read, single-molecule method for mapping protein-DNA interactions genome wide. Nat Methods. 2022;19(6):711–723.

50. Abdulhay NJ, McNally CP, Hsieh LJ, Kasinathan S, Keith A, Estes LS, et al. Massively multiplex single-molecule oligonucleosome footprinting. Elife. 2020;9:e59404.

51. Shipony Z, Marinov GK, Swaffer MP, Sinnott-Armstrong NA, Skotheim JM, Kundaje A, et al. Long-range single-molecule mapping of chromatin accessibility in eukaryotes. Nat Methods. 2020;17(3):319–327.

52. Jeltsch A. On the enzymatic properties of Dnmt1: specificity, processivity, mechanism of linear diffusion and allosteric regulation of the enzyme. Epigenetics. 2006;1(2):63–66.

53. Bestor TH. The DNA methyltransferases of mammals. Hum Mol Genet. 2000;9(16):2395–2402.

54. Goll MG, Bestor TH. Eukaryotic cytosine methyltransferases. Annu Rev Biochem. 2005;74:481–514.

55. Liu Y, Nan B, Niu J, Kapler GM, Gao S. An optimized and versatile counter-flow centrifugal elutriation workflow to obtain synchronized eukaryotic cells. Front Cell Dev Biol. 2021;9:664418.

56. Li H, Handsaker B, Wysoker A, Fennell T, Ruan J, Homer N, et al. The sequence alignment/map format and SAMtools. Bioinformatics. 2009;25(16):2078–2079.

57. Chaisson MJ, Tesler G. Mapping single molecule sequencing reads using basic local alignment with successive refinement (BLASR): application and theory. BMC Bioinform. 2012;13(1):238.

58. Ye J, McGinnis S, Madden TL. BLAST: improvements for better sequence analysis. Nucleic Acids Res. 2006;34(Web Server issue):W6–9.

59. Zhou Y, Fu L, Mochizuki K, Xiong J, Miao W, Wang G. Absolute quantification of chromosome copy numbers in the polyploid macronucleus of Tetrahymena thermophila at the single-cell level. J Eukaryot Microbiol. 2022;69(4):e12907.

60. Seabold, Skipper, Perktold J. Statsmodels: econometric and statistical modeling with python. Proc of the 9th Python in Science Conf. 2010.

61. Quinlan AR, Hall IM. BEDTools: a flexible suite of utilities for comparing genomic features. Bioinformatics. 2010;26(6):841–842.

