## Supplemental file for "Semi-conservative transmission of DNA N^6^-adenine methylation in a unicellular eukaryote"

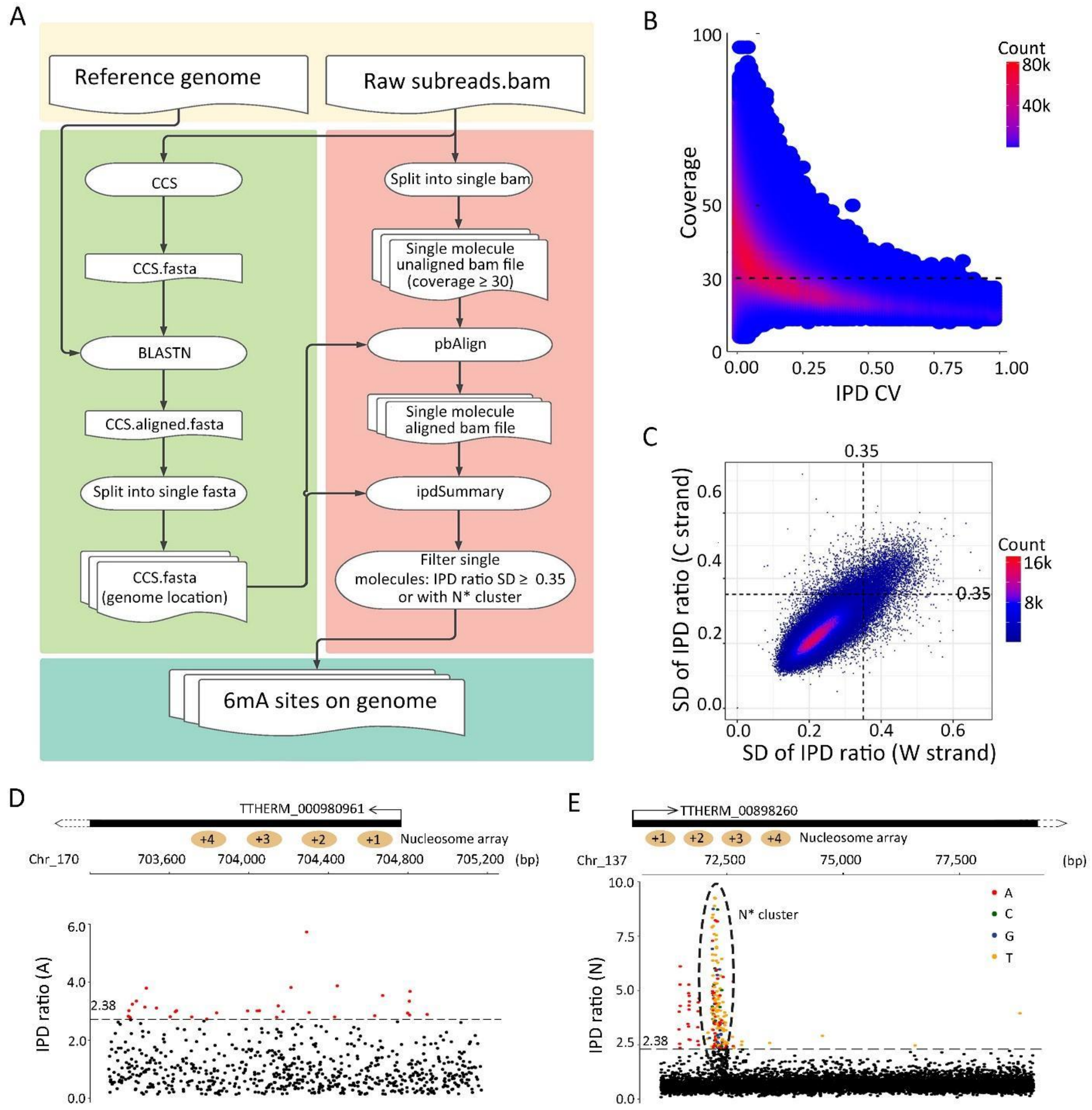


**Figure S1. 6mA detection by SMRT CCS.**

1. 6mA detection by SMRT CCS.

   A. The bioinformatic pipeline for 6mA calling. See methods for details.
   B. Relationship between IPD coefficients of variance (CV=$\frac{tError}{tMean}$, averaged for all adenine sites of a SMRT CCS read) and effective coverage (number of passes for CCS, averaged for all adenine sites in the same read). Read distribution density (count) is plotted as a heat map. A 30× threshold for effective coverage is used to remove reads with noisy IPD values.
   C. Average standard deviation (SD) of IPD ratios for SMRT CCS reads with high effective coverage (≥30×). SD values are calculated for IPD ratios of unmodified adenine sites from W and C, respectively. Read distribution density (count) is plotted as a heat map. The SD threshold (≤0.35), applied to both W and C, is used to remove reads with global anomalies in IPD ratios.
   D. IPD ratios for all A sites in a SMRT CCS read with global anomalies.
   E. IPD ratios for all sites in a SMRT CCS read with local anomalies. Note the high IPD ratio G/C/T sites (as well as A sites) in the N* cluster.

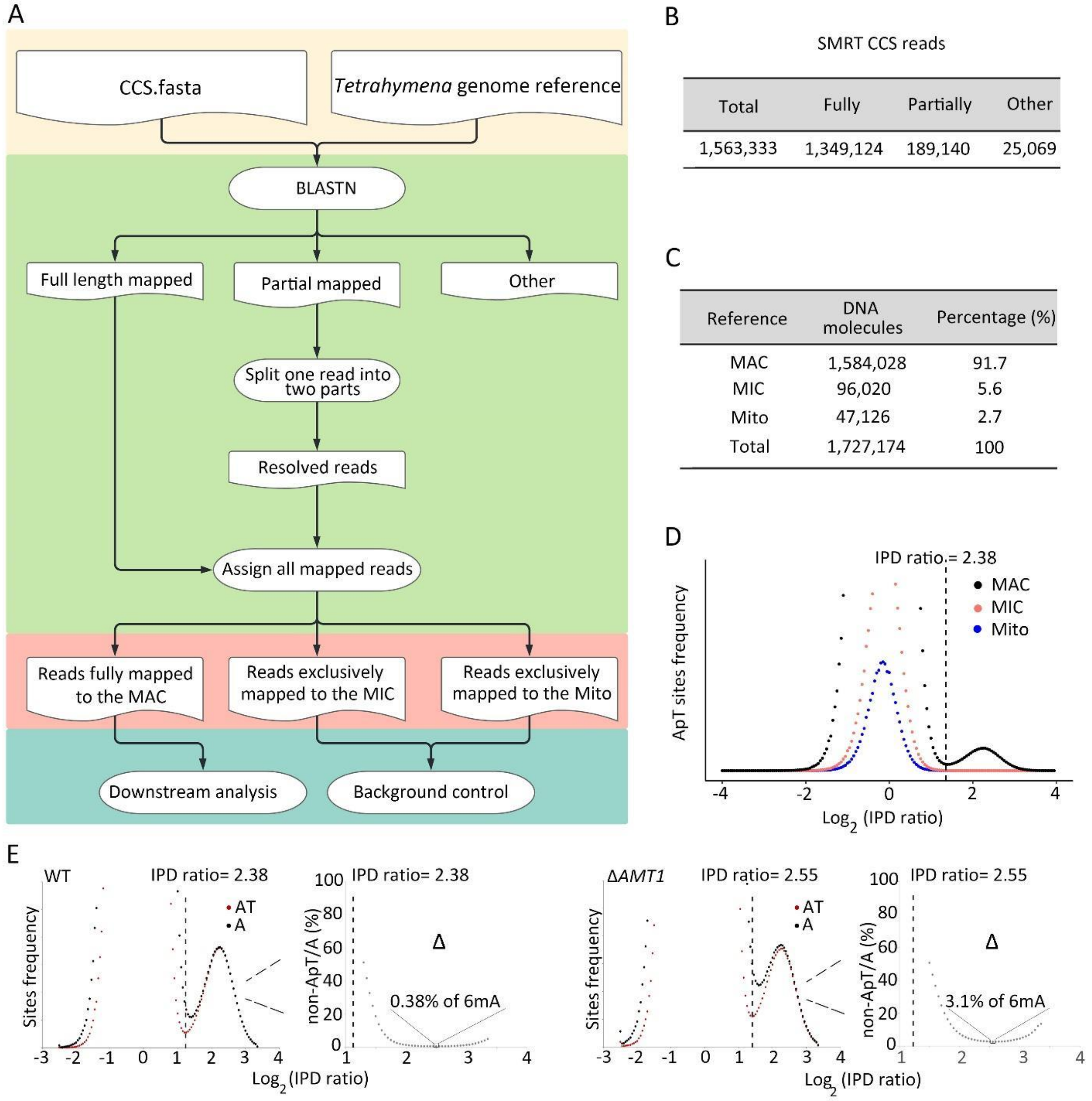


**Figure S2. Comparing 6mA in *Tetrahymena* MAC, MIC, and mitochondrion.**

2. Comparing 6mA in *Tetrahymena* MAC, MIC, and mitochondrion.

   A. The bioinformatic pipeline for mapping CCS reads back to the *Tetrahymena* reference genomes.
   B. Most SMRT CCS reads were either fully or partially mapped to a single locus of *Tetrahymena* reference genomes (MAC, MIC, and mitochondrion) with high confidence. A small percentage of reads were not mapped back with high confidence (Other).
   C. Classification of DNA molecules (after resolving chimeric reads) according to whether they are mapped back to MAC, MIC, or mitochondrion (Mito).
   D. IPD ratio distributions of all A sites in reads mapped back to *Tetrahymena* MAC, MIC, or mitochondrion (Mito). The IPD ratio threshold of 2.38, set for calling 6mApT in MAC, is indicated.
   E. Predominant, if not exclusive, occurrence of 6mA at the ApT dinucleotide in WT (left) and Δ*AMT1* cells (right). IPD ratio distributions for all A sites (black) and A sites at the ApT dinucleotide (red) are plotted. The differential between the two curves (Δ), represented as percentage of non-ApT sites relative to all A sites, is also plotted (only to the right of the 6mA calling threshold); the minimum value of the differential curve is indicated.

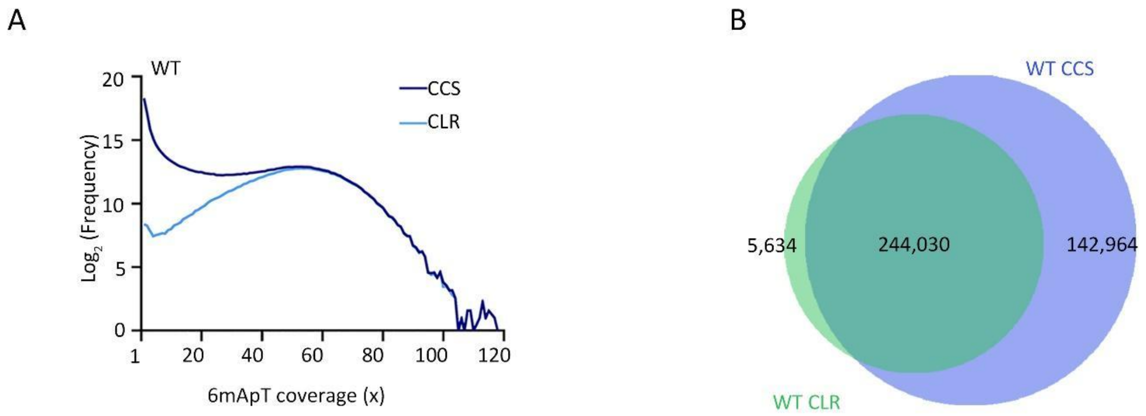


**Figure S3. Comparing SMRT CCS and CLR results for WT *Tetrahymena* cells.**

3. Comparing SMRT CCS and CLR results for WT *Tetrahymena* cells.

   A. CCS and CLR results converge at genomic positions with high 6mApT coverage. For the CCS data (dark blue), x-axis: 6mApT coverage for a genomic position (i.e., the number of reads in which 6mApT has been called in this genomic position); y-axis: number of genomic positions (log2) with the 6mApT coverage indicated in x-axis. We then checked whether 6mApT genomic positions called by CCS were also called by CLR in our previous study (1). For the CLR data (light blue), x-axis: for 6mApT genomic positions called by both CCS and CLR, the 6mApT coverage in CCS; y-axis: number of 6mApT genomic positions (log2) called by both CCS and CLR, with the corresponding 6mApT coverage indicated in x-axis.

   B. Overlap between 6mApT genomic positions called by SMRT CCS (6mApT coverage≥10×) and CLR.

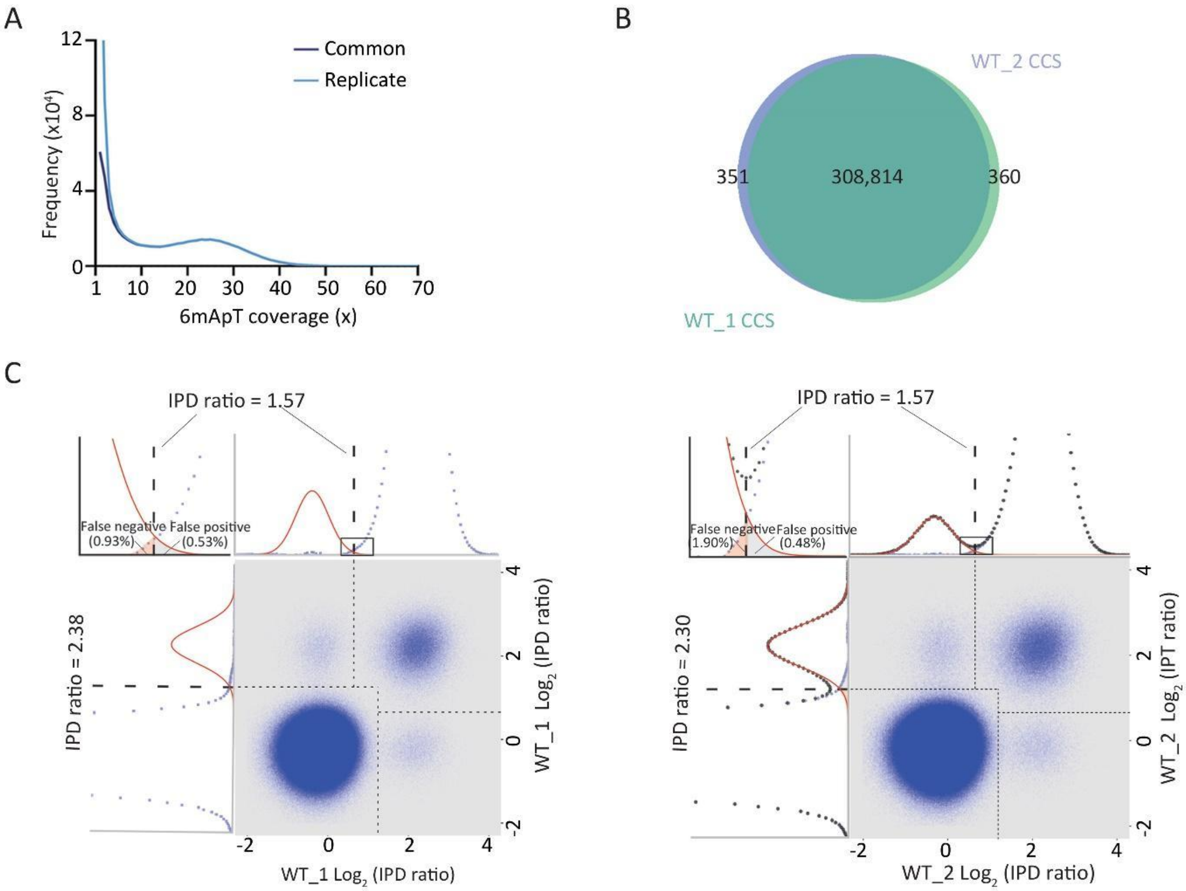


D

| Single molecule | WT_1 | | WT_2 | |
|---|---|---|---|---|
| | Number | Percentage (%) | Number | Percentage (%) |
| ApT | 992,618,784 | 100 | 871,635,758 | 100 |
| 6mApT Sites | 18,750,787 | 1.89 | 17,607,170 | 2.02 |
| full-6mApT | 8,373,723 | 89.32 | 7,872,065 | 89.41 |
| hemi-C-6mApT | 998,137 | 5.32 | 925,862 | 5.25 |
| hemi-W-6mApT | 1,005,024 | 5.36 | 937,178 | 5.32 |
| Total 6mApT | 18,750,787 | 100 | 871,635,758 | 100 |

**Figure S4. Comparing SMRT CCS replicate results for WT *Tetrahymena* cells.**

4. Comparing SMRT CCS replicate results for WT *Tetrahymena* cells.

   A. 6mApT genomic positions called in SMRT CCS replicate results converge at high 6mApT coverage. For one of the two replicate datasets (Replicate: light blue, corresponding to WT_1 dataset), x-axis: 6mApT coverage for a genomic position; y-axis: number of genomic positions with the 6mApT coverage indicated in x-axis. We then checked how many of these 6mApT genomic positions were called in both replicate datasets (Common: dark blue); x-axis: for 6mApT genomic positions called in both datasets, the 6mApT coverage in WT_1 dataset; y-axis: number of 6mApT genomic positions called in both datasets, with the corresponding 6mApT coverage indicated in x-axis.
   B. Overlap between 6mApT genomic positions called in SMRT CCS replicate results (6mApT coverage≥10×).
   C. Demarcation of the four methylation states of ApT duplexes in SMRT CCS replicate results. For details, see Figure 2C: left.
   D. Statistics for 6mApT sites called in SMRT CCS replicate results.

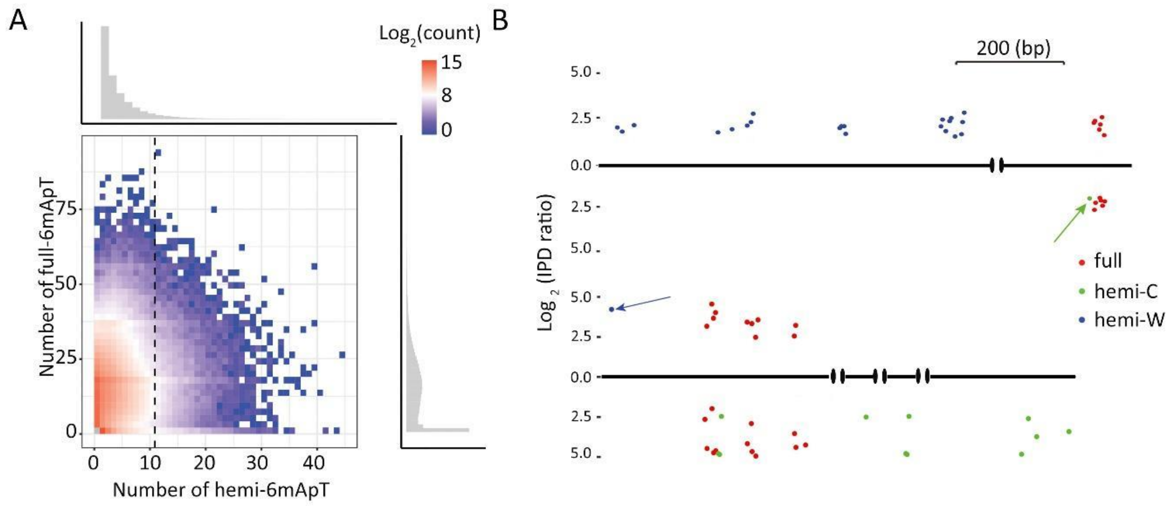


**Figure S5. Hemi⁺ molecules.**

5. Hemi⁺ molecules.

   A. Distribution of CCS reads, according to the number of full-6mApT and hemi-6mApT duplexes they contain. Read distribution density is plotted as a heat map, with the threshold for hemi+ molecules indicated (W+C≥11, dashed line). Read distributions according to separate counts of full-6mApT (right) and hemi-6mApT (top) are also plotted.

   B. Two hemi⁺ molecules with many hemi-6mApT on one strand, but one hemi-6mApT on the other strand (arrow).

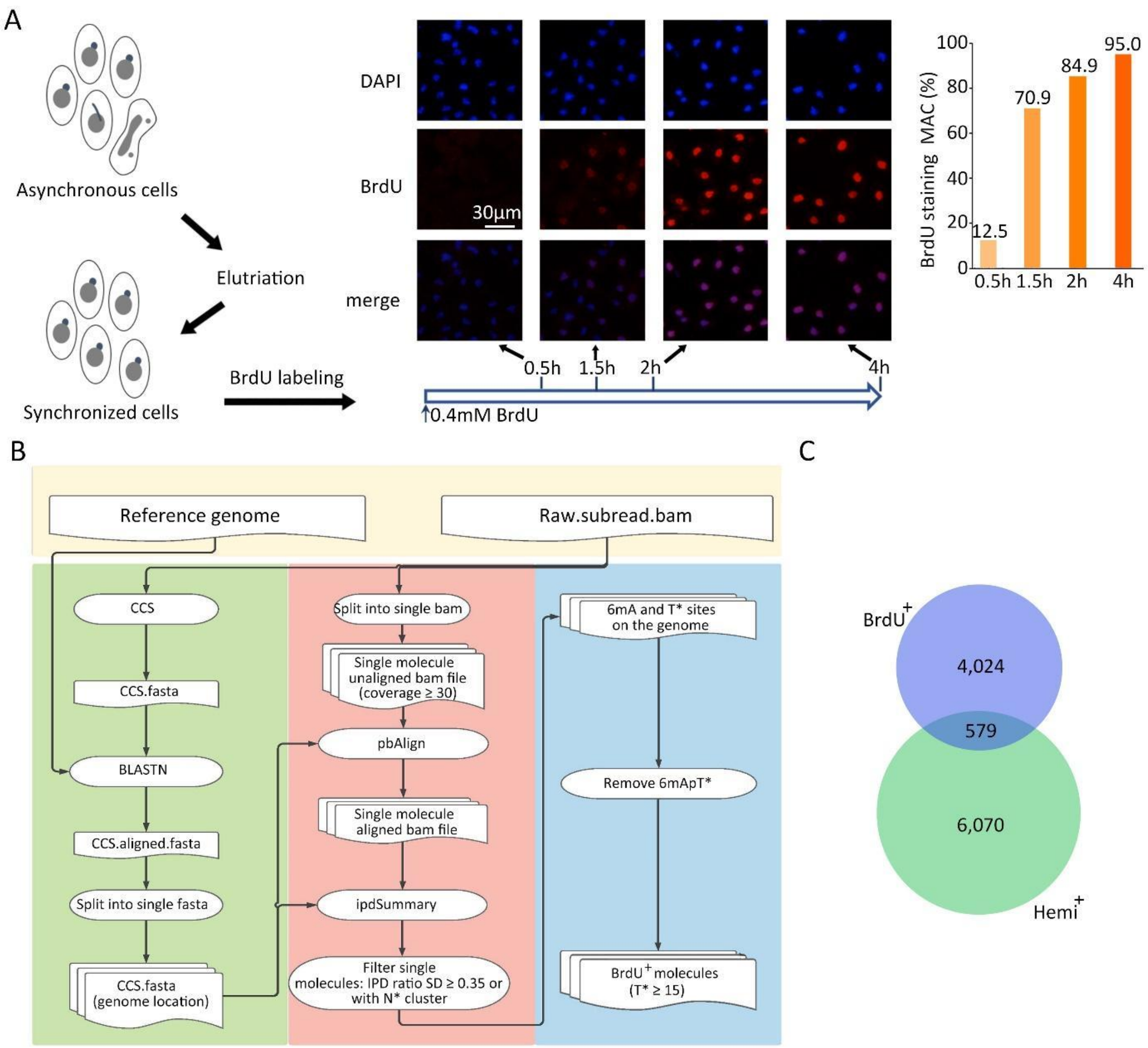


**Figure S6. BrdU$^{+}$ molecules.**

6. BrdU$^+$ molecules.

   A. BrdU-labeling of *Tetrahymena* cells. *Tetrahymena* cells were synchronized at G1 phase by centrifugal elutriation and released for growth in the fresh medium with 0.4mM BrdU (2). *Tetrahymena* samples were taken at 0.5h, 1.5h, 2h, and 4h of labeling. Immunofluorescence (IF) staining showed that the percentage of cells with BrdU signals increased dramatically at 1.5h and plateaued at 2h, indicative of highly synchronous progression through S phase.
   B. The bioinformatic pipeline for calling BrdU$^+$ molecules.
   C. Overlap between hemi$^+$ and BrdU$^+$ molecules, from G1-synchronized cells labeled by BrdU for 1.5h.

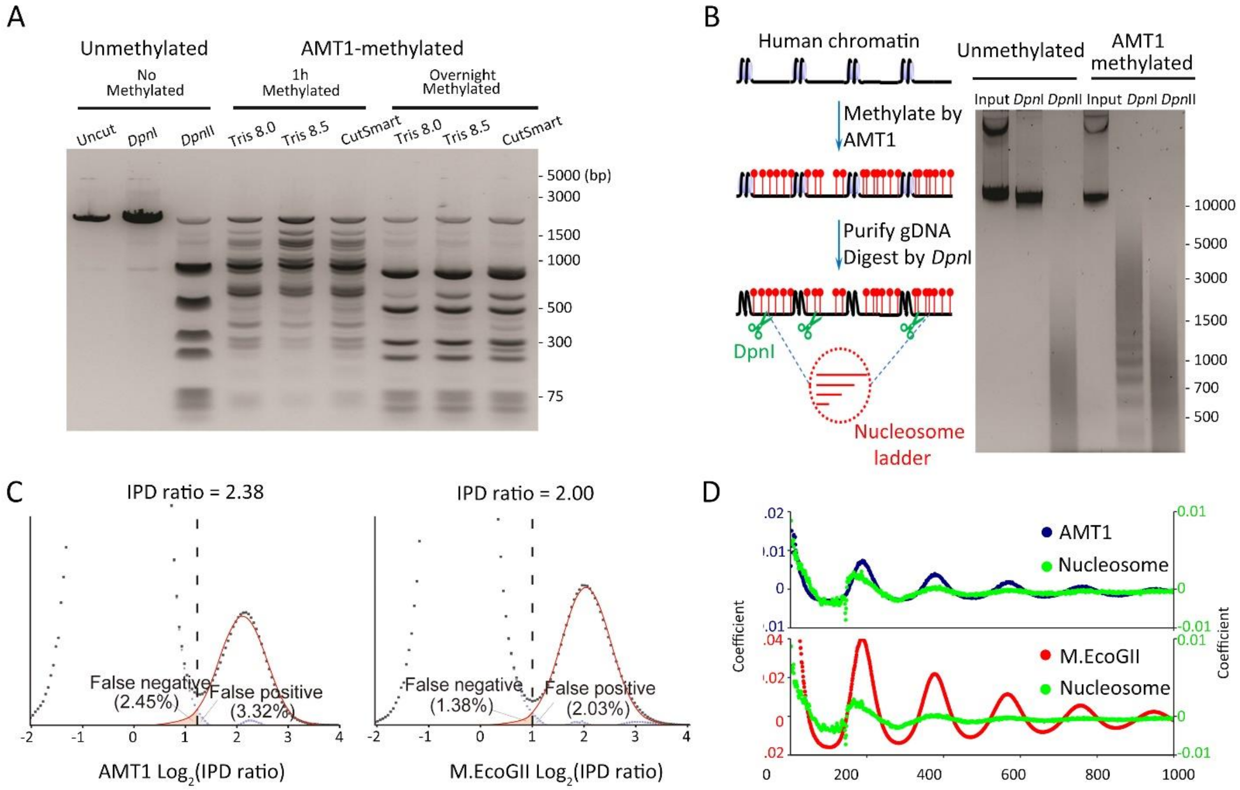


**Figure S7. Additional characterization of *in vitro* methyltransferase activities of AMT1 complex.**

7. Additional characterization of *in vitro* methyltransferase activities of AMT1 complex.

   A. MTase activity of AMT1 complex on linearized plasmid DNA. Three different buffers were tested. Methylation progress was monitored by DpnI cleavage, occurring only at methylated GATC sites. DpnII cleavage, occurring only at unmethylated GATC sites, was used as a control. Note that under all conditions, a substantial fraction of GATC sites were methylated by AMT1 complex in 1h, and almost all overnight, indicative of its robust MTase activity.
   B. MTase activity of AMT1 complex on human chromatin. Methylation progress was monitored by DpnI digestion, with DpnII digestion as a control.
   C. Deconvolution of the 6mA peak and the unmodified A peak for IPD ratio distributions of all A sites, in human chromatin methylated by AMT1 complex (left) or M.EcoGII (right).
   D. Autocorrelation of 6mA distributions at the ensemble level in human chromatin methylated by AMT1 complex and M.EcoGII, respectively. For comparison, autocorrelation of nucleosome distribution in human chromatin was also plotted.

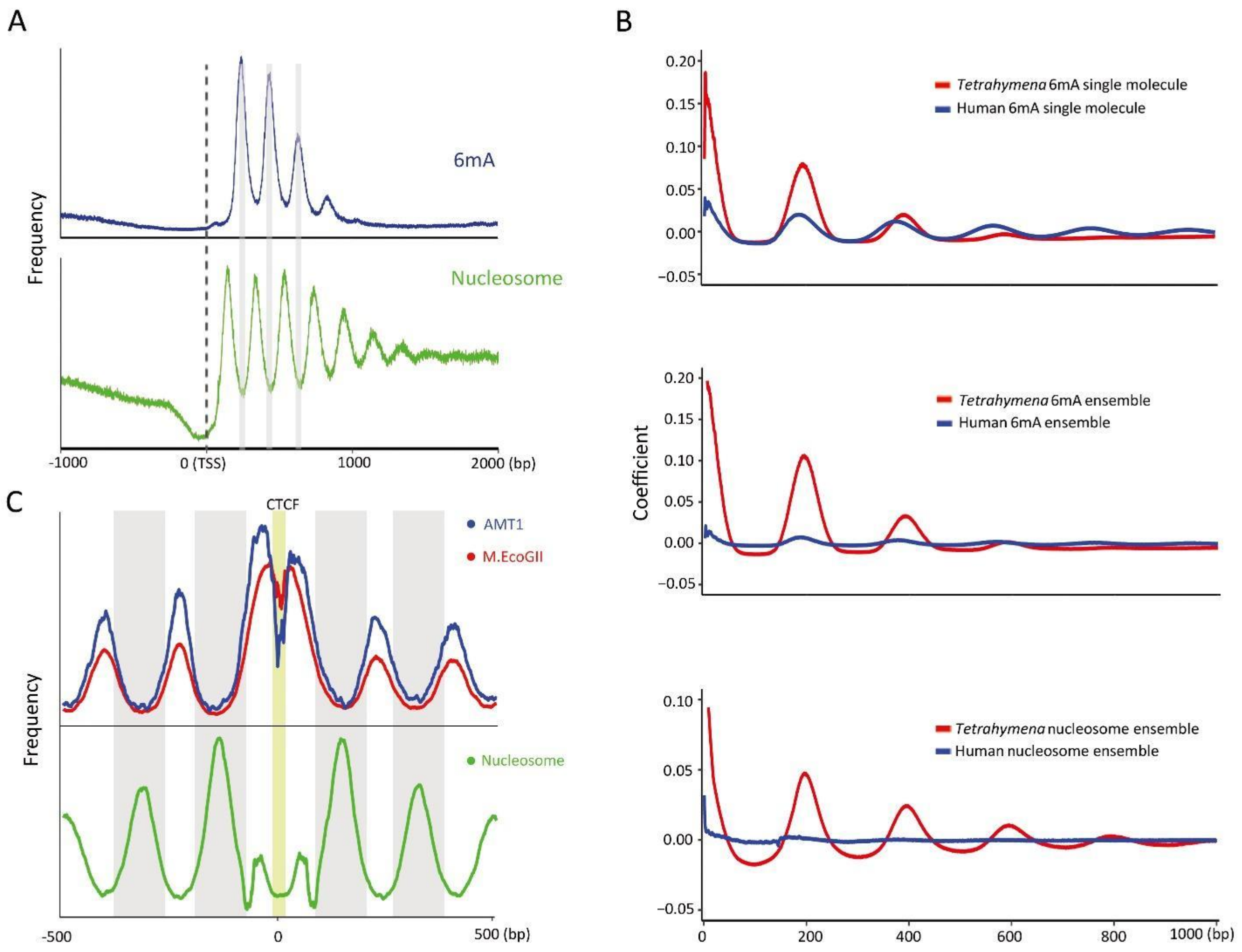


**Figure S8. Additional characterization of the 6mA-nucleosome relationship.**

8. Additional characterization of the 6mA-nucleosome relationship.
   A. 6mA and nucleosome distributions relative to TSS. Pol II-transcribed genes are aligned to TSS; x-axis: the distance downstream (2000 bp) or upstream (-1000 bp) of TSS; y-axis: the aggregated count of 6mA sites (top) or nucleosome dyads (bottom). Note the peak-trough correspondence between the two distributions (gray bars), indicating 6mA enrichment in linker DNA.
   B. Comparing 6mA and nucleosome distributions in *Tetrahymena* cells and *in vitro* methylated human chromatin. From top to bottom: autocorrelation of 6mA distribution at the single molecule level, 6mA distribution at the ensemble level, and nucleosome distribution at the ensemble level.
   C. Anti-correlation between 6mA (top) and nucleosome distributions (bottom) around CTCF-binding sites (yellow highlight) of *in vitro* methylated human chromatin. Note the peak-trough correspondence between the two distributions (gray bars), indicating 6mA enrichment in linker DNA.

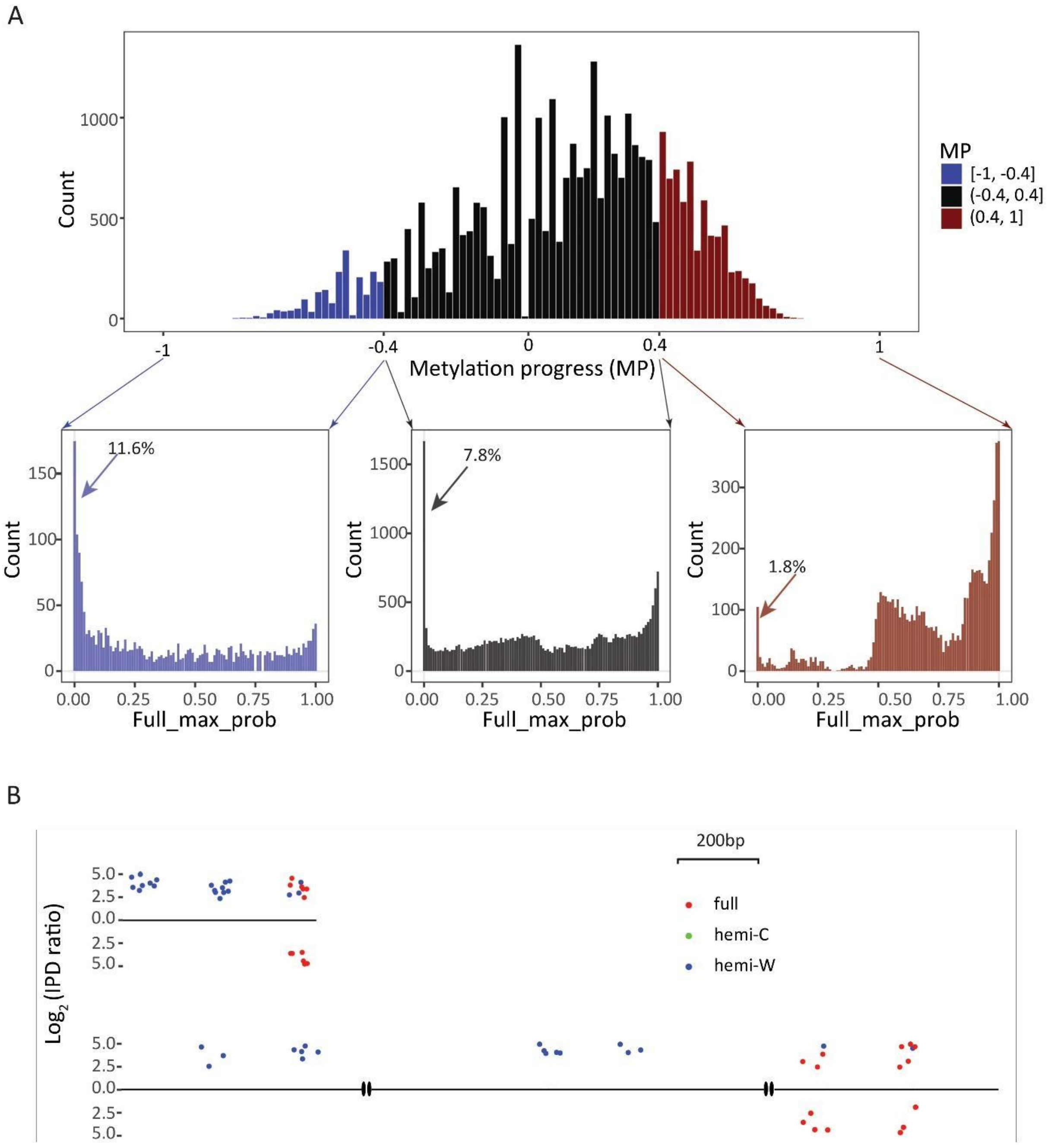


**Figure S9. Full-6mApT congregation in DNA molecules undergoing hemi-to-full conversion in WT *Tetrahymena* cells.**

9. Full-6mApT congregation in DNA molecules undergoing hemi-to-full conversion in WT *Tetrahymena* cells.
    A. Full-6mApT congregation in DNA molecules undergoing hemi-to-full conversion. Methylation progress (MP) is defined for a DNA molecule as the difference-sum ratio between the number of full-6mApT and hemi-6mApT: ($\frac{full-hemi}{full+hemi}$). DNA molecules are divided into three groups according to their MP values (top). Their max inter-full distances are calculated, and their distributions are plotted according to their probabilities in permutated simulations (bottom). The percentage of DNA molecules with probabilities no greater than 0.01 is labeled. Note its decrease with methylation progress.
    B. DNA molecules with congregated full-6mApT intermixed with hemi-6mApT.

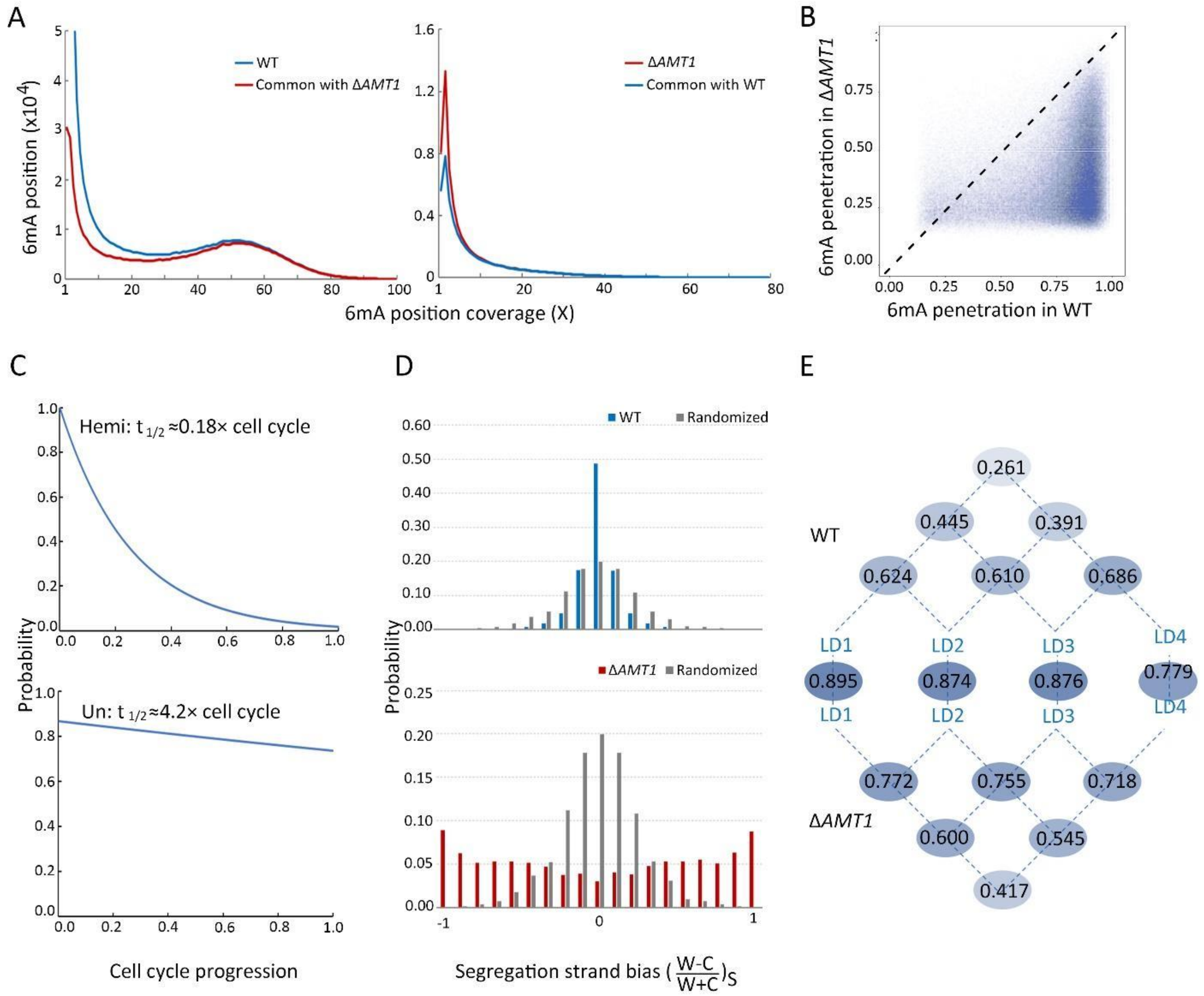


**Figure S10. Comparison of 6mA in WT and Δ*AMT1* cells.**

10. Comparison of 6mA in WT and Δ*AMT1* cells.

A. Overlap between methylated ApT positions in the MAC genome of WT and Δ*AMT1* cells. Note their convergence with increasing 6mA coverage.

B. Comparing 6mA penetration of individual genomic positions present in both WT and Δ*AMT1* cells. Minimal 6mA coverage: 10. Note that most positions are below the diagonal line, indicative of reduced 6mA penetration in Δ*AMT1* cells.

C. Exponential decay kinetics for hemi-6mApT in WT cells, and unmethylated ApT (at methylatable positions) in Δ*AMT1* cells.

D. 6mA segregation strand bias. In Δ*AMT1* cells, DNA molecules exhibited much stronger 6mA segregation strand bias than the randomized control (bottom), due to slow *de novo* methylation. In WT cells, DNA molecules exhibited even less 6mA segregation strand bias than the randomized control (top), due to quick restoration of full methylation. Note that both hemi and full-6mApT were counted here (hemi once for the corresponding strand; full twice, one for W and one for C). This is different from calculating segregation strand bias for $hemi^+$ molecules, in which only hemi-6mApT was counted.

E. 6mA levels of individual linker DNAs (LDs) in the gene body in WT and Δ*AMT1* cells are strongly correlated. For each LD, 6mA level is quantified by sum of penetration values for all methylated ApT positions within. LD1 is defined to be in between the +1 and +2 nucleosome (the first and second nucleosome downstream of TSS, belong to the canonical nucleosome array in the gene body); LD2-4 are defined iteratively further downstream. Spearman's rank correlation coefficients were calculated for all possible pairs between LD1-4 within WT and Δ*AMT1* cells, respectively. They were also calculated between equivalent LDs from WT and Δ*AMT1* cells.

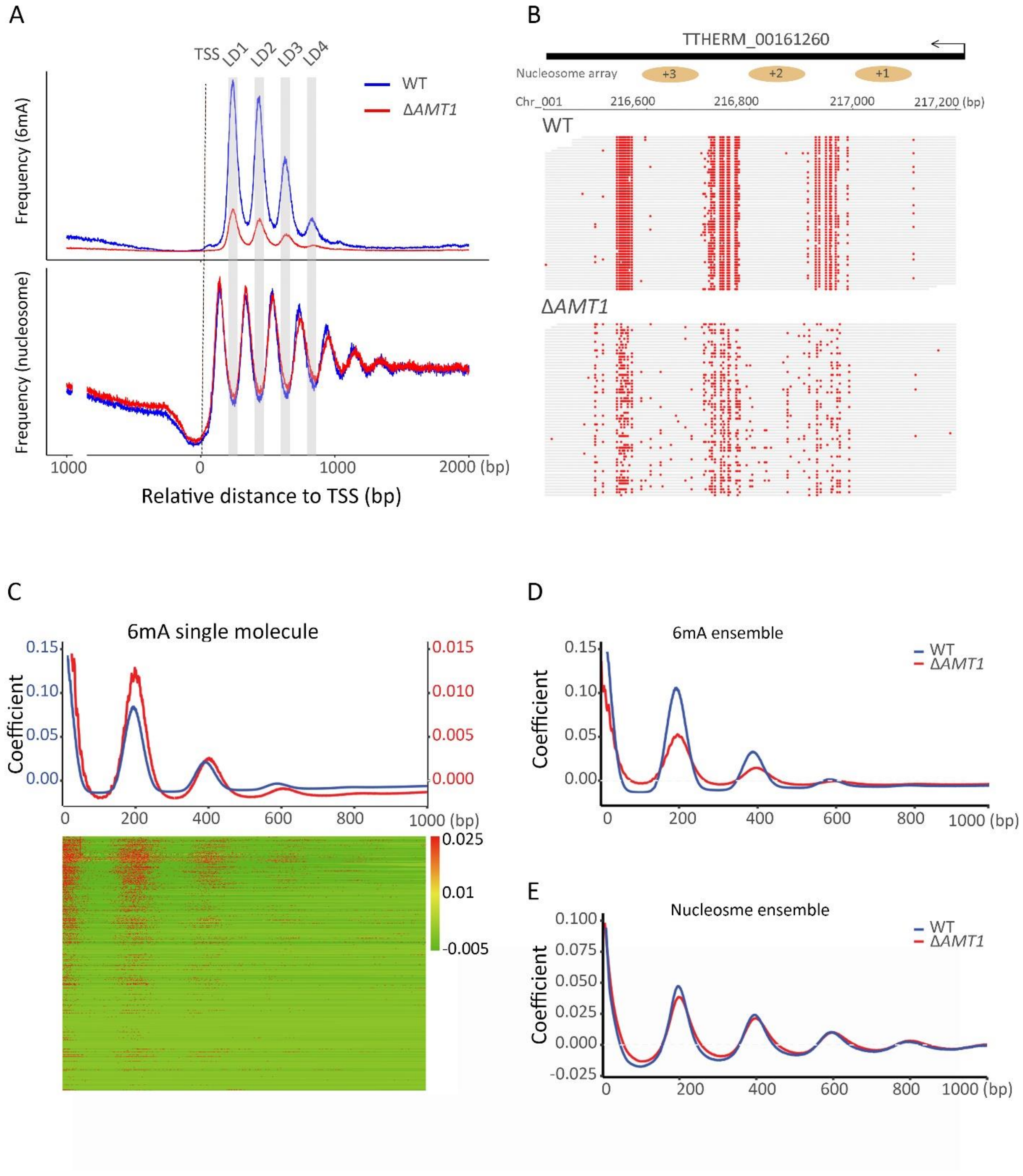


**Figure S11. Dispersion of 6mA in Δ*AMT1* cells.**

11. Dispersion of 6mA in Δ*AMT1* cells.

   A. 6mA and nucleosome distributions along the gene body in WT and Δ*AMT1* cells. Pol II-transcribed genes are aligned to TSS; x-axis: distance upstream (-1000bp) or downstream (2000bp) of TSS; y-axis: cumulative counts of 6mA sites (top) and nucleosome dyads (bottom).
   B. 6mApT sites in DNA molecules from WT (top) and Δ*AMT1* cells (bottom). Note 6mA dispersion in the latter.
   C. Periodic 6mA distribution at the single molecule level in Δ*AMT1* cells. Autocorrelation between 6mA sites (distance ≤1kb) was calculated for individual DNA molecules, ranked by their median absolute deviations, and plotted as heat maps (bottom) and aggregated correlograms (top, along with the WT curve).
   D. Periodic 6mA distributions at the ensemble level in WT and Δ*AMT1* cells.
   E. Periodic nucleosome distributions at the ensemble level in WT and Δ*AMT1* cells.

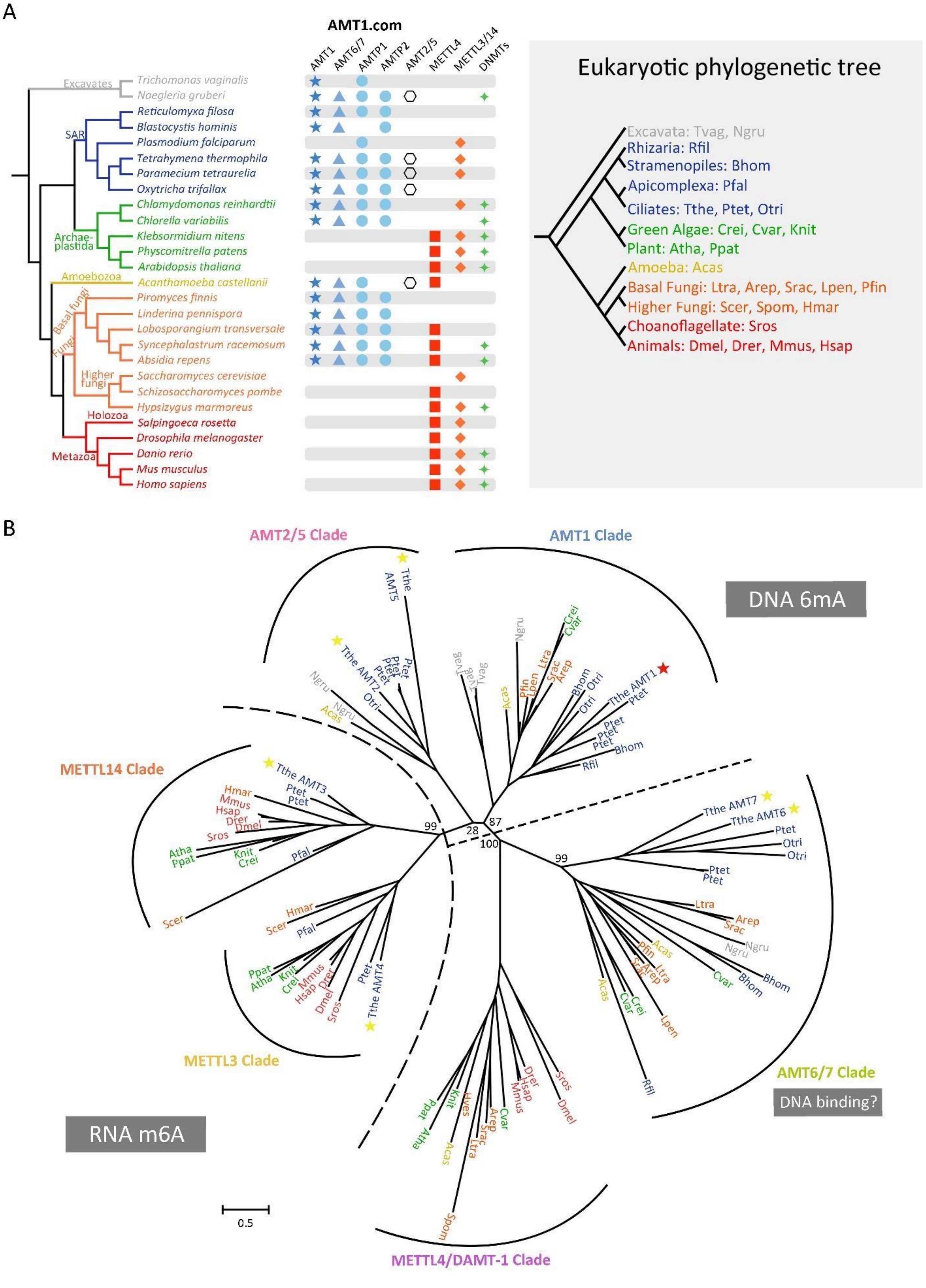

A
AMT1.com
AMT1
AMT6/7
AMTP1
AMTP2
AMT2/5
METTL4
METTL3/14
DNMTs
Excavates
Trichomonas vaginalis
Naegleria gruberi
SAR
Reticulomyxa filosa
Blastocystis hominis
Plasmodium falciparum
Tetrahymena thermophila
Paramecium tetraurelia
Oxytricha trifallax
Archae-plastida
Chlamydomonas reinhardtii
Chlorella variabilis
Klebsormidium nitens
Physcomitrella patens
Arabidopsis thaliana
Amoebozoa
Acanthamoeba castellanii
Basal fungi
Fungi
Piromyces finnis
Linderina pennispora
Lobosporangium transversale
Syncephalastrum racemosum
Absidia repens
Higher fungi
Saccharomyces cerevisiae
Schizosaccharomyces pombe
Hypsizygus marmoreus
Holozoa
Salpingoeca rosetta
Drosophila melanogaster
Metazoa
Danio rerio
Mus musculus
Homo sapiens
Eukaryotic phylogenetic tree
Excavata: Tvag, Ngru
Rhizaria: Rfil
Stramenopiles: Bhom
Apicomplexa: Pfal
Ciliates: Tthe, Ptet, Otri
Green Algae: Crei, Cvar, Knit
Plant: Atha, Ppat
Amoeba: Acas
Basal Fungi: Ltra, Arep, Srac, Lpen, Pfin
Higher Fungi: Scer, Spom, Hmar
Choanoflagellate: Sros
Animals: Dmel, Drer, Mmus, Hsap
B
AMT2/5 Clade
AMT1 Clade
DNA 6mA
Tthe AMT5
Tthe AMT2
Tthe AMT1
METTL14 Clade
Tthe AMT3
Tthe AMT7
Tthe AMT6
99
28
87
100
99
99
METTL3 Clade
Tthe AMT4
AMT6/7 Clade
DNA binding?
RNA m6A
0.5
METTL4/DAMT-1 Clade

**Figure S12. Phylogenetic distribution of MT-A70 MTases in eukaryotes.**

12. Phylogenetic distribution of MT-A70 MTases in eukaryotes.

A. Left: distribution of MT-A70 family MTases (for 6mA and m6A) and DNMT family MTases (for 5mC) in main eukaryote groups. Right: major branches of eukaryotic evolution.
B. Phylogenetic tree of MT-A70 MTases. Note that members in AMT1 and AMT2/5 clades generally contain the DPPW motif critical for catalysis, while members in AMT6/7 and METTL4/DAMT-1 clade lack it. AMT7 (and possibly AMT6) is the heterodimeric partner for AMT1, likely providing a target recognition domain for binding the double-stranded DNA substrate. This situation is analogous to the heterodimeric RNA m6A MTase METTL3 and METTL14 (3,4). Plants, fungi, and animals have only members of METTL4/DAMT-1 clade, but not AMT1, AMT2/5, and AMT6/7 clades. METTL4/DAMT-1 clade members therein are more likely to be involved in DNA binding than 6mA deposition.

| Reference | 6mApT/ApT (%) | 6mApA/ApA (%) | 6mApC/ApC (%) | 6mApG/ApG (%) |
|---|---|---|---|---|
| MAC | 1.89 | 0.027 | 0.059 | 0.081 |
| MIC | 0.017 | 0.021 | 0.061 | 0.090 |
| Mito | 0.014 | 0.014 | 0.054 | 0.070 |

**Table S1. 6mA levels in MAC, MIC, or mitochondrion.**

Percentage of 6mApT/ApA/ApC/ApG sites called in MAC, MIC, or mitochondrion (based on the IPD ratio threshold of 2.38), relative to all ApT/ApA/ApC/ApG sites.

### Supplementary Methods

#### *In vitro* methylation of plasmid DNA

1μg of pUC19 plasmid generated from *dam-/dcm-* competent *E. coli* (NEB) was linearized by EcoRI and methylated with AMT1 complex in a 20μL reaction (20mM Tris-HCl (pH 8.0 or 8.5) or 1×CutSmart Buffer (NEB), 160μM SAM, 2mM EDTA, 0.5mM EGTA). Reactions were performed for 1h at 37°C or overnight at 30°C, quenched with addition of 1% SDS, and incubated with proteinase K for 1h at 50°C. Methylated plasmid was digested with DpnI. Digestion products of methylated samples and the unmethylated control were resolved on 1% agarose gel.

#### SMRT CCS data analysis

6mA sites are associated with significantly increased IPD ratios; at sufficient abundance, 6mA sites form a peak distinct from unmodified adenines in the IPD ratio distribution. We deconvoluted the 6mA peak and the unmodified A peak of the ApT dinucleotide by fitting the former with a Gaussian distribution. We set the 6mA calling threshold at the intersection of the two peaks (IPD ratio=2.38, for WT *Tetrahymena*), which keeps both false positive and false negative rates at low levels (FP: 1.93%, FN: 1.12%). For the same dataset, if one increases the threshold (e.g., IPD ratio=3.03), then FP will decrease while FN will increase (FP: 0.90%, FN: 10.15%). Furthermore, as the threshold is determined by the relative abundance of 6mA and unmodified A, it exhibits variation across different datasets (higher threshold for lower 6mA abundance). This is also relevant for distinguishing hemi-6mApT and full-6mApT in WT *Tetrahymena*. For an ApT site reverse complementary to a 6mApT site in ApT duplexes, the unmodified A peak was instead dominated by the 6mA peak; this caused the 6mA calling threshold to shift substantially to the left (IPD ratio=1.57). Similar analyses were performed for genomic DNA samples from Δ*AMT1* cells and Δ*MLL1* cells, as well as the human chromatin *in vitro* methylated by AMT1 complex and M.EcoGII.

We aligned CCS reads to the latest *Tetrahymena* genome references for the MAC (5), MIC (unpublished, based on the published MIC genome (6), with improved assembly of repetitive sequences), and mitochondrion (7). Discrepancy from the MAC reference mostly originated from heterogeneous junctions of MAC-destined sequences, generated by imprecise removal of thousands of MIC-limited sequences. Discrepancy from the MIC reference was mostly found in repetitive regions that are difficult to assemble. The mapping results of representative genomic regions were visualized using Jbrowse (8) or ggplot2 package in R (9).

The SMRT CCS raw data and 6mA calling results can be downloaded from:

https://dataview.ncbi.nlm.nih.gov/object/PRJNA932808?reviewer=tak7a6fv74pn85n6nmi3ekpdqb

The code for base modification calling is available at:

https://gitfront.io/r/user-1129035/86c191cfcd32dc94d0047fa0617509701f714764/Pacbio_m6A_calling/

### CCS versus CLR

SMRT CCS allowed us to call 6mA on individual DNA molecules. Many genomic positions were heterogeneously represented as either 6mA or unmodified A sites on different DNA molecules. 6mA at these positions would have been more difficult to detect by SMRT sequencing at the ensemble level (CLR), due to mixing of 6mA and unmodified A signals. Indeed, comparison between the CCS and CLR results showed that many more genomic positions with 6mA were detected by CCS, especially for genomic positions with low 6mA penetration.

### Gene level analyses

Models for well-annotated Pol II-transcribed genes in *Tetrahymena* (15,810 in total) were updated with the latest RNA-seq data (SRR9176464). We calculated the coefficients of variance (CV) for 6mA levels across different DNA molecules covering a gene. We only counted DNA molecules fully covering the gene body. For comparison, we only included genes with high coverage (≥20× overall coverage) and high 6mA levels (ΣP≥2) in both

WT and Δ*AMT1* cells. CV ratios between WT and Δ*AMT1* cells were calculated as an indicator of their relative 6mA variability at the gene level.

For composite analysis, *Tetrahymena* genes were aligned to TSS and TES (the gene body is normalized to unit length) and extended in both directions by 0. 5kb. Alternatively, *Tetrahymena* genes are aligned to TSS with upstream (1 kb) and downstream (2 kb) extensions. For 6mA and nucleosome distributions, counts of 6mA sites and MNase-seq fragment centers in specified genomic regions were aggregated.

#### 6mA and nucleosome distributions around CTCF binding sites

CTCF ChIP-seq data for a human leukemia cell line OCI-AML3 were used to call CTCF peaks with MACS2 (10). CTCF motif profile (accession number: MA0139.1) was downloaded from JASPER (11) (http://jaspar.genereg.net/) and sed to scan all the CTCF peak regions by FIMO (12). High quality CTCF binding sites (motif score ≥ 60) were retained for further analyses. For 6mA distribution, 6mA sites were aggregated for each base in a window of -500 to 500 bp around the high quality CTCF binding sites (aligned to the left boundary of the CTCF binding consensus sequence). For nucleosome distribution, Micro-C data were downloaded from the 4DN data portal (https://data.4dnucleome.org) and processed as previously described (13). The raw data adapters were trimmed with Trim Galore and mapped with Bowtie2 (14) against the human genome (hg19) using the following parameters (bowtie2 --local -q --phred33 --threads 12 -x hg19 bowtie2 indexed genome -U trimmed.fq.gz -S output.sam). Multiple mapped reads and reads with low mapping quality (MAPQ < 30) were removed using samtools (15). PCR duplicates were removed with Picard tools (http://broadinstitute.github.io/picard/). Read starts were then shifted by 73 bp to reveal positions of nucleosome dyads, which were aggregated for each base in a window of -500 to 500 bp around the high quality CTCF binding sites.

#### Phylogenetic analysis

The well-studied 27 taxa covered major branches of eukaryotic evolution were selected to search for homologous proteins (16-35). List of species: *Naegleria gruberi*,

*Trichomonas vaginalis*, *Reticulomyxa filosa*, *Blastocystis hominis*, *Plasmodium falciparum*, *Paramecium tetraurelia*, *Tetrahymena thermophila*, *Oxytricha trifallax*, *Chlamydomonas reinhardtii*, *Chlorella variabilis*, *Klebsormidium nitens*, *Arabidopsis thaliana*, *Physcomitrella patens*, *Acanthamoeba castellanii* , *Lobosporangium transversale*, *Absidia repens*, *Syncephalastrum racemosum*, *Linderina pennispora*, *Piromyces finnis*, *Saccharomyces cerevisiae*, *Schizosaccharomyces pombe*, *Hypsizygus marmoreus*, *Salpingoeca rosetta*, *Drosophila melanogaster*, *Danio rerio*, *Mus musculus*, and *Homo sapiens*. The AMTs 1–7, AMTP1 and AMTP2 amino acid sequences were queried against the database using PSI-BLAST (36) (maximum E-value = 1e–4), respectively. Retrieved hits were collapsed using CD-HIT (37) (-c 0.97) to remove redundant sequences. Sequences were aligned using MUSCLE (38) and phylogenetic trees were constructed using FastTree (39) under default parameters.

### References


1. Wang, Y., Sheng, Y., Liu, Y., Zhang, W., Cheng, T., Duan, L., Pan, B., Qiao, Y., Liu, Y. and Gao, S. A distinct class of eukaryotic MT-A70 methyltransferases maintain symmetric DNA $N^6$-adenine methylation at the ApT dinucleotides as an epigenetic mark associated with transcription. Nucleic Acids Res. 2019;47:11771-11789.
2. Liu, Y., Nan, B., Niu, J., Kapler, G.M. and Gao, S. An optimized and versatile counter-flow centrifugal elutriation workflow to obtain synchronized eukaryotic cells. Front Cell Dev Biol. 2021;9:664418.
3. Wang, X., Feng, J., Xue, Y., Guan, Z., Zhang, D., Liu, Z., Gong, Z., Wang, Q., Huang, J., Tang, C. et al. Structural basis of N(6)-adenosine methylation by the METTL3-METTL14 complex. Nature. 2016;534:575-578.
4. Wang, P., Doxtader, K.A. and Nam, Y. Structural basis for cooperative function of mettl3 and mettl14 methyltransferases. Mol Cell. 2016;63:306-317.
5. Sheng, Y., Duan, L., Cheng, T., Qiao, Y., Stover, N.A. and Gao, S. The completed macronuclear genome of a model ciliate *Tetrahymena thermophila* and its application in genome scrambling and copy number analyses. Sci China Life Sci. 2020;63:1534-1542.
6. Hamilton, E.P., Kapusta, A., Huvos, P.E., Bidwell, S.L., Zafar, N., Tang, H., Hadjithomas, M., Krishnakumar, V., Badger, J.H., Caler, E.V. et al. Structure of the germline genome of *Tetrahymena thermophila* and relationship to the massively rearranged somatic genome. Elife. 2016;5: e19090.
7. Brunk, C.F., Lee, L.C., Tran, A.B. and Li, J. (2003) Complete sequence of the mitochondrial genome of *Tetrahymena thermophila* and comparative methods for identifying highly divergent genes. Nucleic Acids Research, 31, 1673-1682.
8. Skinner, M.E., Uzilov, A.V., Stein, L.D., Mungall, C.J. and Holmes, I.H. JBrowse: A next-generation genome browser. Genome Res. 2009;19:1630-1638.
9. Wickham, H. (2016) ggplot2: elegant graphics for data analysis. O' Reilly Press, USA.
10. Zhang, Y., Liu, T., Meyer, C.A., Eeckhoute, J., Johnson, D.S., Bernstein, B.E., Nusbaum, C., Myers, R.M., Brown, M. and Li, W. Model-based analysis of ChIP-Seq (MACS). Genome Bio. 2008;9:1-9.
11. Fornes, O., Castro-Mondragon, J.A., Khan, A., Van der Lee, R., Zhang, X., Richmond, P.A., Modi, B.P., Correard, S., Gheorghe, M. and Baranašić, D. JASPAR 2020: update of the open-access database of transcription factor binding profiles. Nucleic Acids Res. 2020;48:D87-D92.
12. Grant, C.E., Bailey, T.L. and Noble, W.S. FIMO: scanning for occurrences of a given motif. Bioinformatics. 2011;27:1017-1018.
13. Krietenstein, N., Abraham, S., Venev, S.V., Abdennur, N., Gibcus, J., Hsieh, T.S., Parsi, K.M., Yang, L., Maehr, R., Mirny, L.A. et al. Ultrastructural details of mammalian chromosome architecture. Mol Cell. 2020;78:554-565.
14. Ramírez, F., Ryan, D.P., Grüning, B., Bhardwaj, V., Kilpert, F., Richter, A.S., Heyne, S., Dündar, F. and Manke, T. deepTools2: a next generation web server for deep-sequencing data analysis. Nucleic Acids Res. 2016;44:W160-W165.
15. Li, H., Handsaker, B., Wysoker, A., Fennell, T., Ruan, J., Homer, N., Marth, G., Abecasis, G. and Durbin, R. The sequence alignment/map format and SAMtools. Bioinformatics. 2009;25:2078-2079.

16. Fritz-Laylin, L.K., Prochnik, S.E., Ginger, M.L., Dacks, J.B., Carpenter, M.L., Field, M.C., Kuo, A., Paredez, A., Chapman, J., Pham, J. et al. The genome of *Naegleria gruberi* illuminates early eukaryotic versatility. Cell. 2010;140:631-642.
17. Aurrecoechea, C., Brestelli, J., Brunk, B.P., Carlton, J.M., Dommer, J., Fischer, S., Gajria, B., Gao, X., Gingle, A., Grant, G. et al. GiardiaDB and TrichDB: integrated genomic resources for the eukaryotic protist pathogens *Giardia lamblia* and *Trichomonas vaginalis*. Nucleic Acids Res. 2009;37:D526-D530.
18. Glöckner, G., Hülsmann, N., Schleicher, M., Noegel, Angelika A., Eichinger, L., Gallinger, C., Pawlowski, J., Sierra, R., Euteneuer, U., Pillet, L. et al. The Genome of the Foraminiferan *Reticulomyxa filosa*. Curr Biol. 2014;24:11-18.
19. Denoeud, F., Roussel, M., Noel, B., Wawrzyniak, I., Da Silva, C., Diogon, M., Viscogliosi, E., Brochier-Armanet, C., Couloux, A., Poulain, J. et al. Genome sequence of the stramenopile Blastocystis, a human anaerobic parasite. Genome Bio. 2011;12:R29.
20. Aurrecoechea, C., Brestelli, J., Brunk, B.P., Dommer, J., Fischer, S., Gajria, B., Gao, X., Gingle, A., Grant, G., Harb, O.S. et al. PlasmoDB: a functional genomic database for malaria parasites. Nucleic Acids Res. 2009;37:D539-D543.
21. Arnaiz, O., Meyer, E. and Sperling, L. (2020) ParameciumDB 2019: integrating genomic data across the genus for functional and evolutionary biology. Nucleic Acids Res. 2020;48:D599-D605.
22. Stover, N.A., Krieger, C.J., Binkley, G., Dong, Q., Fisk, D.G., Nash, R., Sethuraman, A., Weng, S. and Cherry, J.M. Tetrahymena Genome Database (TGD): a new genomic resource for *Tetrahymena thermophila* research. Nucleic Acids Res. 2006;34:D500-D503.
23. Swart, E.C., Bracht, J.R., Magrini, V., Minx, P., Chen, X., Zhou, Y., Khurana, J.S., Goldman, A.D., Nowacki, M., Schotanus, K. et al. (2013) The *Oxytricha trifallax* macronuclear genome: a complex eukaryotic genome with 16,000 tiny chromosomes. PLoS Biol. 2013;11:e1001473.
24. Merchant, S.S., Prochnik, S.E., Vallon, O., Harris, E.H., Karpowicz, S.J., Witman, G.B., Terry, A., Salamov, A., Fritz-Laylin, L.K., Maréchal-Drouard, L. et al. The *Chlamydomonas* genome reveals the evolution of key animal and plant functions. Science. 2007;318:245-250.
25. Berardini, T.Z., Reiser, L., Li, D., Mezheritsky, Y., Muller, R., Strait, E. and Huala, E. The arabidopsis information resource: Making and mining the “gold standard” annotated reference plant genome. Genesis. 2015;53:474-485.
26. Rensing, S.A., Lang, D., Zimmer, A.D., Terry, A., Salamov, A., Shapiro, H., Nishiyama, T., Perroud, P.F., Lindquist, E.A., Kamisugi, Y. et al. The Physcomitrella genome reveals evolutionary insights into the conquest of land by plants. Science. 2008;319:64-69.
27. Aurrecoechea, C., Barreto, A., Brestelli, J., Brunk, B.P., Caler, E.V., Fischer, S., Gajria, B., Gao, X., Gingle, A., Grant, G. et al. AmoebaDB and MicrosporidiaDB: functional genomic resources for Amoebozoa and Microsporidia species. Nucleic Acids Res. 2011;39:D612-D619.
28. Mondo, S.J., Dannebaum, R.O., Kuo, R.C., Louie, K.B., Bewick, A.J., LaButti, K., Haridas, S., Kuo, A., Salamov, A., Ahrendt, S.R. et al. Widespread adenine $N^6$-methylation of active genes in fungi. Nat Genet. 2017;49:964-968.

29. Cherry, J.M., Hong, E.L., Amundsen, C., Balakrishnan, R., Binkley, G., Chan, E.T., Christie, K.R., Costanzo, M.C., Dwight, S.S., Engel, S.R. et al. Saccharomyces Genome Database: the genomics resource of budding yeast. Nucleic Acids Res. 2012;40:D700-D705.
30. Lock, A., Rutherford, K., Harris, M.A., Hayles, J., Oliver, S.G., Bähler, J. and Wood, V. (2019) PomBase 2018: user-driven reimplementation of the fission yeast database provides rapid and intuitive access to diverse, interconnected information. Nucleic Acids Res. 2019;47:D821-D827.
31. Harris, T.W., Arnaboldi, V., Cain, S., Chan, J., Chen, W.J., Cho, J., Davis, P., Gao, S., Grove, C.A., Kishore, R. et al. WormBase: a modern model organism information resource. Nucleic Acids Res. 2020;48:D762-D767.
32. Larkin, A., Marygold, S.J., Antonazzo, G., Attrill, H., dos Santos, G., Garapati, P.V., Goodman, Joshua L., Gramates, L S., Millburn, G., Strelets, V.B. et al. FlyBase: updates to the *Drosophila melanogaster* knowledge base. Nucleic Acids Res. 2021;49: D899-D907.
33. Karimi, K., Fortriede, J.D., Lotay, V.S., Burns, K.A., Wang, D.Z., Fisher, M.E., Pells, T.J., James-Zorn, C., Wang, Y., Ponferrada, V G. et al. Xenbase: a genomic, epigenomic and transcriptomic model organism database. Nucleic Acids Res. 2018;46:D861-D868.
34. Ruzicka, L., Howe, D.G., Ramachandran, S., Toro, S., Van Slyke, C.E., Bradford, Y.M., Eagle, A., Fashena, D., Frazer, K., Kalita, P. et al. The Zebrafish Information Network: new support for non-coding genes, richer Gene Ontology annotations and the Alliance of Genome Resources. Nucleic Acids Res. 2019;47:D867-D873.
35. Bult, C.J., Blake, J.A., Smith, C.L., Kadin, J.A., Richardson, J.E. and the Mouse Genome Database, G. Mouse Genome Database (MGD) 2019. Nucleic Acids Res. 2019;47:D801-D806.
36. Altschul, S.F., Madden, T.L., Schäffer, A.A., Zhang, J., Zhang, Z., Miller, W. and Lipman, D.J. Gapped BLAST and PSI-BLAST: a new generation of protein database search programs. Nucleic Acids Res. 1997;25:3389-3402.
37. Li, W. and Godzik, A. Cd-hit: a fast program for clustering and comparing large sets of protein or nucleotide sequences. Bioinformatics. 2006;22:1658-1659.
38. Edgar, R.C. MUSCLE: a multiple sequence alignment method with reduced time and space complexity. BMC Bioinformatics. 2004;5:113.
39. Price, M.N., Dehal, P.S. and Arkin, A.P. FastTree 2 – approximately maximum-likelihood trees for large alignments. PLoS ONE. 2010;5:e9490.
